# The phagocytosis oxidase/Bem1p (PB1) domain-containing protein PB1CP negatively regulates the NADPH oxidase RBOHD in plant immunity

**DOI:** 10.1101/2020.12.28.423414

**Authors:** Yukihisa Goto, Noriko Maki, Jan Sklenar, Paul Derbyshire, Frank L.H. Menke, Cyril Zipfel, Yasuhiro Kadota, Ken Shirasu

## Abstract

- Perception of pathogen-associated molecular patterns (PAMPs) by surface-localized pattern-recognition receptors activates RESPIRATORY BURST OXIDASE HOMOLOG D (RBOHD) through direct phosphorylation by BOTRYTIS-INDUCED KINASE 1 (BIK1) and induces the production of reactive oxygen species (ROS). ROS have direct antimicrobial properties but also serve as signaling molecules to activate additional defense responses such as stomatal closure. RBOHD activity must be tightly controlled to avoid the detrimental effects of ROS, but little is known about RBOHD downregulation.
- To better understand the regulation of RBOHD, we used co-immunoprecipitation of RBOHD coupled with mass spectrometry analysis to identify RBOHD-associated proteins.
- Among RBOHD-associated proteins, we identified PHAGOCYTOSIS OXIDASE/ BEM1P (PB1) DOMAIN-CONTAINING PROTEIN (PB1CP). We found that PB1CP negatively regulates RBOHD and the resistance against the fungal pathogen *Colletotrichum higginsianum*. PB1CP directly interacts with RBOHD *in vitro*, and PAMP treatment increases the interaction *in vivo*. PB1CP is localized at the cell periphery and in cytoplasm, but PAMP treatment induces PB1CP relocalization to small endomembrane compartments. *PB1CP* overexpression reduces plasma membrane localization of RBOHD, suggesting that PB1CP down-regulates RBOHD function by relocalizing it away from the plasma membrane.
- We reveal a novel negative regulation mechanism of ROS production through the relocalization of RBOHD by PB1CP.

## Introduction

The production of reactive oxygen species (ROS) is an immune response against infection that is well-conserved across biological kingdoms. ROS not only have antimicrobial activities, but they also act as signaling molecules to induce additional immune responses (Lambeth *et al*., 2000). Excessive ROS production can have detrimental effects on cellular functions by damaging DNA, proteins, lipids, and other macromolecules (Lorrain, 2003; Moeder & Yoshioka, 2008). As such, ROS production must be produced in the right amount and place, and at the right time to minimize cellular damage.

NADPH oxidases (NOXs) are highly conserved plasma- and endo-membrane enzymes that play a crucial role in ROS production in plants, animals, and fungi (Segal, 2016). NOXs transfer electrons from cytosolic NADPH or NADH to apoplastic oxygen, leading to the production of superoxide (O_2_^-^). O_2_^-^ can then be converted to hydrogen peroxide (H_2_O_2_) by superoxide dismutases (Suzuki *et al*., 2011; Marino *et al*., 2012; Kadota *et al*., 2015). In animals, the activity of NOX proteins is tightly controlled by regulatory proteins and Ca^2+^. The 91-kD glycoprotein subunit of phagocyte oxidase (GP91^phox^), also known as NADPH oxidase 2 (NOX2), is the best-characterized NOX. NOX2 forms a heterodimer with the membrane protein p22^phox^ and together they bind to the cytosolic regulators p47^phox^, p67^phox^, p40^phox^, and the small GTPase Rac. Interaction with these regulators leads to the activation of NOX2 (Canton & Grinstein, 2014). Mutations in NOX2 or its regulatory proteins cause chronic granulomatous disease in which patients suffer from chronic or recurrent bacterial and fungal infections due to the absence of an oxygen burst (Bedard & Krause 2007), thus showing the crucial role that NOXs play in immunity. In contrast to NOX2, NOX5 and DUOX have additional EF-hand motifs at the N-terminal domain (ND), suggesting their regulation by Ca^2+^-binding (Canton & Grinstein 2014). In addition, calcium-dependent kinases, such as protein kinase Cα and Ca^2+^/calmodulin-dependent protein kinase II, are known to phosphorylate and activate NOX5 and DUOX.

In plants, NOXs belong to the respiratory burst oxidase homolog (RBOH) family, which contains ten members in *Arabidopsis thaliana* (Torres & Dangl 2005; Kadota *et al*., 2015). Among RBOHs, RBOHD plays a particularly crucial role in immunity and stress responses. RBOHD-produced ROS are required for the induction of numerous defense responses, including callose deposition, stomatal closure, and systemic acquired resistance (Mishina & Zeier, 2007; Suzuki *et al*., 2013; Ross *et al*., 2014; Mittler & Blumwald, 2015). Recent studies have clarified the RBOHD activation mechanisms that are triggered after the perception of pathogen-associated molecular patterns (PAMPs) by surface-localized pattern recognition receptors (PRRs). Activated RBOHD induces PAMP-induced ROS production as one of the readouts in so-called pattern-triggered immunity (PTI) (Kadota *et al*., 2014; Li *et al*., 2014). Leucine-rich repeat receptor kinases (LRR-RKs) EFR and FLS2, which are the PRRs for the immunogenic peptides of bacterial EF-Tu or flagellin (elf18 or flg22), respectively, induce instantaneous association with the coreceptor LRR-RK BAK1. The PRR complex interacts directly with, and phosphorylates receptor-like cytoplasmic kinases (RLCKs) such as BOTRYTIS INDUCED KINASE 1 (BIK1). RBOHD forms a complex with EFR and FLS2, and phosphorylated BIK1 interacts directly with, and phosphorylates specific residues at the N-terminal domain (ND) of RBOHD, which is required for RBOHD activation (Kadota *et al*., 2014; Li *et al*., 2014). In addition to the regulation by RLCKs, Ca^2+^-based regulation is also required for RBOHD activation. PAMP perception by PRRs activates plasma membrane Ca^2+^ channels such as OSCA1.3 and the cyclic nucleotide-gated channel (CNGC) proteins CNGC2 and CNGC4 (at least under specific Ca^2+^ concentrations) through BIK1, which lead to the influx of Ca^2+^ (Tian *et al*., 2019; Thor *et al*., 2020). Ca^2+^ in turn activates RBOHD through Ca^2+^ binding to the EF-hand motif in RBOHD-ND as well as through phosphorylation by Ca^2+^-dependent protein kinases (CPKs). Like NOX2, small GTPase Rac binds to the ND of the RBOHD homolog in rice, which is important for ROS production (Wong *et al*., 2007). Although RBOHs and NOX2 are structurally similar, none of the known NOX2 regulators has a RBOH regulatory function in plants, except for RAC.

How RBOHD activity is down-regulated is not well understood. Endocytosis and vacuolar degradation are known to decrease membrane proteins and their activity of the downstream signaling pathways. For example, many PRRs are endocytosed after ligand perception through the clathrin-mediated pathway (Robatzek *et al*., 2006; Mbengue *et al*., 2016). Similar to PRRs, RBOHD is also endocytosed under salt stress (Hao *et al*., 2014), suggesting the involvement of endocytosis in down-regulating RBOHD activity. Interestingly, PBL13, an RLCK, phosphorylates the C-terminal domain (CD) of RBOHD, which triggers the ubiquitination of RBOHD by PIRE (PBL13-INTERACTING RING DOMAIN E3 LIGASE) (Lee *et al*., 2020a). Because PIRE-mediated ubiquitination of RBOHD lowers RBOHD levels, ubiquitination may serve as a signal for endocytosis and vacuolar degradation. However, it is not clear whether endocytosis or vacuolar degradation of RBOHD are induced as a mechanism to halt PAMP-induced ROS production.

In this work, we identify phagocytosis oxidase/Bem1p (PB1) domain-containing protein (PB1CP) as a novel negative regulator of RBOHD. The PB1 domain is conserved in animal NOX2 regulators p40^phox^ and p67^phox^, and plays important roles for their assembly with NOX2, as well as its activation, suggesting that there is cross-kingdom conservation of functional domains among NOX regulators. In contrast to the positive regulation of p40^phox^ and p67^phox^, PB1CP negatively regulates PAMP-induced ROS production as well as resistance against the fungal pathogen *Colletotrichum higginsianum*. PB1CP directly interacts with RBOHD and competes with BIK1 for binding. PAMP treatment induces direct interaction between PB1CP and RBOHD, suggesting that PB1CP specifically interacts with the activated form of RBOHD. PB1CP localizes at the cell periphery and the cytoplasm, but PAMP treatment results in the accumulation of PB1CP within small endomembrane compartments. In addition, *PB1CP* overexpression reduces the plasma membrane localization of RBOHD, suggesting that PB1CP relocalizes RBOHD from the plasma membrane to small endomembrane compartments. Co-treatment with PAMP and cycloheximide (CHX), a protein synthesis inhibitor, results in localization of RBOHD to dot-like PB1CP endomembrane compartments. These data suggest a novel negative regulation mechanism of ROS production through PB1CP-mediated relocalization of RBOHD.

## Materials and Methods

### Plant materials and growth conditions

Seeds of Arabidopsis (*Arabidopsis thaliana*) ecotype Columbia (Col-0), T-DNA insertion mutant of *pb1cp-1* (SALK_036544), and *pb1cp-2* (SALK_207053) were sown on soil or half-strength Murashige-Skoog media containing 1 % sucrose. After two days of cold treatment to break seed dormancy, plants were grown under short-day photoperiods (8 h light/16 h dark) at 23 °C. Liquid culture seedlings were grown under continuous light at 23 °C. T-DNA insertion lines were genotyped using primers listed in Table S3.

### Vector construction and generation of transgenic lines

The CDS region of *PB1CP* was amplified by PCR with KoD FX neo (Toyobo, Japan), and the resulting PCR product was cloned into the epiGreenB5 (3×HA) and epiGreenB(eGFP) vectors between the *Cla*I and *Bam*HI restriction sites with an In-Fusion HD Cloning Kit (Clontech, USA) to generate epiGreenB5-Cauliflower mosaic virus (CaMV) *p35S::PB1CP-3*×*HA* and epiGreenB-CaMV *p35S::PB1CP-3-eGFP* for transient expression assays in *Nicotiana benthamiana* and for stable transformation of Arabidopsis (Nekrasov *et al*., 2009). All of the other genes encoding candidate RBOHD-associated proteins were cloned into epiGreenB5 by the same strategy using an In-Fusion HD Cloning Kit (Clontech, USA). To generate epiGreenB-*pPB1CP::PB1CP-3-eGFP* for transient expression in *N. benthamiana*, amplicon containing the 2000-bp promoter upstream of start codon and coding regions of *PB1CP* was cloned into epiGreenB (eGFP) between the *Eco*RI and *Bam*HI restriction sites (Nekrasov *et al*., 2009). To generate epiGreenB-CaMV *p35S::mScarlet-RBOHD*, the coding regions of *RBOHD* and mScarlet were amplified by PCR and cloned into the epiGreenB5 between *Cla*I and *Bam*HI restriction sites. Arabidopsis stable transgenic lines of *p35S::PB1CP-3*×*HA* were generated based on the floral drop (Clough & Bent, 1998) or floral dip (Martinez-Trujillo *et al*., 2004) methods as described previously. PCR primers for these constructs are listed in Table S3.

### Protein extraction and co-immunoprecipitation

Protein extraction and immunoprecipitation were performed as described previously (Kadota *et al*., 2014, 2016) with minor modifications. Ten grams fresh weight of seedlings were used for large-scale immunoprecipitation to identify RBOHD-associated proteins, and two grams were used for directed co-immunoprecipitations. After seedlings were ground in liquid nitrogen with sand (Sigma-Aldrich), extraction buffer (50 mM Tris-HCl, pH 7.5, 150 mM NaCl, 10 % glycerol, 5 mM DTT, 2.5 mM NaF, 1 mM Na_2_MoO_4_•2H_2_O, 0.5 % [w/v] polyvinylpyrrolidone, 1 % [v/v] P9599 Protease Inhibitor Cocktail (Sigma-Aldrich), 100 μM phenylmethylsulphonyl fluoride and 2 % [v/v] IGEPAL CA-630 (Sigma-Aldrich), 2 mM EDTA and 1 % [v/v] protein phosphatase inhibitor cocktail 2 and 3 (Sigma-Aldrich)) were added at a concentration of 2 mL/g tissue powder. Samples were incubated at 4 °C for 1 h and clarified by several centrifugations at 13,000 rpm for 20 min at 4 °C. Supernatant protein concentrations were adjusted to 5 mg/mL and incubated for 2 h at 4 °C with 200 μL of anti-FLAG matrix (Sigma-Aldrich) for co-immunoprecipitation with FLAG-RBOHD and with 200 μL of anti-HA magnetic beads (Miltenyi Biotec) for co-immunoprecipitation with PB1CP-HA. Beads were then washed three times with extraction buffer. FLAG peptides were used for the elution of FLAG-RBOHD to eliminate non-specific interactions with the beads. PB1CP-HA was eluted with boiled sodium dodecyl sulfate (SDS) sample buffer.

### Protein identification by LC-MS/MS

The identification of proteins by LC-MS/MS was performed as previously described (Ntoukakis *et al*., 2009; Kadota *et al*., 2014). In brief, proteins were separated by SDS-PAGE (NuPAGe®, Invitrogen) and after staining with Coomassie Brilliant Blue (CBB) (SimplyBlue™ stain, Invitrogen), the proteins were excised from the gel and digested with trypsin. LC-MS/MS analysis was performed using a LTQ-Orbitrap mass-spectrometer (Thermo Scientific) and a nanoflow-HPLC system (nanoAcquity; Waters) as described previously (Ntoukakis *et al*., 2009). The TAIR10 database (www.Arabidopsis.org) using the Mascot algorithm (Matrix Science). The Scaffold program (Proteome Software) was used to validate MS/MS-based peptide and protein identifications and to annotate spectra.

### Transient expression in *N. benthamiana*

*Agrobacterium tumefaciens* AGL1 strains carrying the binary expression vectors were grown in LB medium supplemented with the appropriate antibiotics. Cultures were pelleted by centrifugation and re-suspended in buffer containing 10 mM MgCl_2_, 10 mM MES pH 5.6, and 100 μM acetosyringone to a concentration with OD_600_ = 0.3 and incubated for 3 h at room temperature. *Agrobacterium* strains were syringe-infiltrated into the same leaf.

### ROS burst assay

A ROS burst assay was performed as described previously (Kadota *et al*., 2014). ROS production in *N. benthamiana* was induced with 1 μM flg22, and measured using 8 leaf discs three days after Agroinfiltration. Candidate-associated proteins of RBOHD and GUS were expressed in the same leaves, and ROS production was compared to measure the effect of the candidate on flg22-induced ROS production. For ROS burst assay in Arabidopsis, eight leaf discs from four-to six-week-old soil-grown plants (three plants per genotype, 24 leaf discs in total) were used. Discs were punched out of leaves using a cork borer, then floated overnight on sterile water. The water was replaced with a reaction solution containing the chemiluminescent probe Luminol (Wako, Japan) at 400 nM or L-012 (Wako, Japan) at 1 μM, horseradish peroxidase (HRP) at 20 g/ml (Sigma-Aldrich, USA), and one of the PAMPs (1 μM flg22, 1 μM elf18, or 10 μM chitin). Luminescence was measured over 30 min with a Tristar^2^ multimode reader (Berthold Technologies, Germany).

### Quantitative RT-PCR (RT-qPCR)

RT-qPCR was performed as described previously (Kadota *et al*., 2019). Total RNA was extracted from 2-week-old Arabidopsis seedlings using an RNeasy Plant Mini Kit (Qiagen, Germany) according to the manufacturer’s instructions. RNA was reverse transcribed with a ReverTraAce qPCR RT Kit (Toyobo, Japan) according to the manufacturer’s instructions. One μg of total RNA was used as a template for cDNA synthesis. RT-qPCR was carried out using Thunderbird SYBR qPCR Mix (Toyobo, Japan) with a Stratagene mx 3000p real-time thermal cycler (Agilent, USA). Relative transcript levels were calculated against a standard curve with normalization to the expression of *PLANT U-BOX PROTEIN1* (*PUB1*) gene (*AT5G15400*, Azevedo *et al*., 2001). Primers used for the RT-qPCR are listed in Table S3.

### MAPK activation assay

MAPK activation assays were performed as described previously (Goto *et al*., 2020). Six-week-old Arabidopsis seedling samples were flash-frozen with liquid nitrogen, and proteins were extracted in protein extraction buffer (50 mM Tris-HCl pH 7.5, 150 mM NaCl, 10 % glycerol, 2 mM EDTA, 5 mM DTT, 1 × EDTA-free Complete Protease Inhibitor Cocktail [Roche, USA], 0.1 % IGEPAL CA630, 0.5 mM PMSF, 1 mM Na_2_MoO_4_, 1 mM NaF, 0.5 mM Na_3_VO_4_, 20 mM β-glycerophosphate). The extract was centrifuged at 16000 × g to remove insoluble material, and supernatant protein concentration was measured using the Bradford method (Bio-Rad Laboratories, USA). Total proteins were separated by SDS-PAGE and blotted onto a PVDF membrane as recommended by the manufacturer (Transblot, Bio-Rad Laboratories, USA). The membrane was blocked overnight at 4 °C in a solution of 5 % (w/v) skim milk (Wako, Japan) in Tris-buffered saline with 0.05 % (v/v) Tween 20 (TBS-T). Phosphorylated MAPKs were detected using α-phospho-p44/42 MAPK (Erk1/2) (Thr202/Tyr204) (D13.14.4E) rabbit monoclonal antibody (1:2000, Cell Signaling Technology, USA) for 1 h at room temperature in a solution of 5 % (w/v) BSA (Sigma-Aldrich, Japan) in TBS-T, followed by incubation with α-rabbit IgG-HRP-conjugated secondary antibodies (1:10000, Roche, USA) for 1 h at room temperature in a solution of 5 % skim milk in TBS-T. HRP-conjugated antibody signal was detected using Super Signal West Femto Maximum Sensitivity Substrate (Thermo Fisher Scientific, USA) with a LAS 4000 system (GE Healthcare, USA). The PVDF membranes were stained with CBB to verify equal loading.

### Pathogen infection assay

Bacterial infection assays were performed using *Pseudomonas syringae* pv. *tomato* (*Pto*) DC3000 *COR*^*-*^ and *P. syringae* pv. *cilantro* (*Pci*) 0788-9 as described previously (Zipfel *et al*., 2004; Goto *et al*., 2020). Bacterial strains were grown overnight in LB medium containing 500 μg/mL kanamycin and 100 μg/mL rifampicin. Cells were harvested by centrifugation, and pellets were re-suspended in 10 mM MgCl_2_ to 1.0 × 10^6^ CFU/ml. Immediately prior to spraying, 0.02 % (v/v) Silwet L-77 was added to bacterial suspensions, and bacteria were sprayed until saturation onto leaf surfaces of 5-to 6-week-old plants. Leaf discs were taken three days post-inoculation from three leaves per plant and six plants per genotype. Leaf discs were ground in 10 mM MgCl_2_, diluted, and plated on LB agar with appropriate selection. Plates were incubated at 28 °C and colonies were counted two days later.

*C. higginsianum* infection assays were performed as described previously (Hiruma & Saijo, 2016a,b; Goto *et al*., 2020). Conidia were harvested from 7-to 10-day-old potato dextrose agar cultures under 12 h of near-UV blue light and 12 h of darkness at 25 °C. Conidia were washed three times with distilled water, and counted using a Bright-line Hemocytometer (Hausser Scientific, USA). Five droplets, each containing 5.0×10^6^ conidia/mL, were spotted on leaves of 3-week-old Arabidopsis plants. Infected plants were watered and maintained in a covered tray. At 7-to 10-days post-infection, macroscopic symptoms were assessed by measuring lesions along their X- and Y-axes using Fiji software (Schindelin *et al*., 2012).

### *In vitro* pull-down assay

Ten micrograms of MBP and GST fusion proteins were incubated in pull-down buffer (20 mM HEPES-KOH pH 7.5, 50 mM KCl, 5 mM MgCl_2_, 1 % Tween 20, 1 mM DTT, and 100 μM phenylmethylsulphonyl fluoride) at 4 °C for 1 h. MBP and GST fusion proteins were separated from supernatants using amylose resin (New England Biolabs) and Glutathione Sepharose 4 Fast Flow (GE Healthcare Life Sciences), respectively. Amylose resin and Glutathione Sepharose were washed four times with pull-down buffer and eluted with 10 mM maltose and 10 mM reduced glutathione, respectively. Bound proteins were visualized by SDS-PAGE/CBB staining.

### Confocal microscopy

Four-week-old *N. benthamiana* leaves were used to observe subcellular localization of PB1CP-GFP and mScarlet-RBOHD. The fluorescence signals of GFP and mScarlet were recorded using confocal laser scanning microscopy (Leica TCS SP5, Leica Microsystems GmbH, Germany) after excitation at 488 nm or 561 nm, respectively with an argon laser. Fluorescence emission was collected between 500 − 540 nm for GFP and 559 − 595 nm for mScarlet. Stacking images of 30 consecutive 1μm planes were displayed as a maximum projection. The micrographs were processed using LAS X version; 3.3.0.16799 and Fiji software (Schindelin *et al*., 2012).

## Results

### PB1CP is a novel interactor of NADPH Oxidase RBOHD during PTI

To understand the regulatory mechanism of PAMP-induced ROS production during PTI, we employed co-immunoprecipitation coupled with liquid chromatography-tandem mass spectrometry (LC-MS/MS) to identify novel regulators of RBOHD in Arabidopsis. We used a stable transgenic Arabidopsis line expressing 3×FLAG-tagged RBOHD under the control of its own promoter in a *rbohD* knockout background (*rbohD*/*pRBOHD::3*×*FLAG-gRBOHD*) (Kadota *et al*., 2014). Plants were treated with elf18 or with elf18 and flg22 simultaneously (elf18+flg22) to activate RBOHD. FLAG-RBOHD was immunoprecipitated using α-FLAG antibody and eluted by competition with FLAG peptide, and RBOHD-associated proteins were identified by LC-MS/MS (Fig. **S1a**; Table S1). In three independent experiments, we identified 450 candidate RBOHD-associated proteins. Among those candidates there were proteins known to associate with RBOHD, such as cysteine-rich RK 2 (CRK2) (Kimura *et al*., 2020), and extra-large guanine nucleotide-binding protein 3 (XLG3) (Liang *et al*., 2016). There were also known PRR interactors such as BAK1(Chinchilla *et al*., 2007; Heese *et al*., 2007), CERK1 (Miya *et al*., 2007), FER (Stegmann *et al*., 2017), IOS1(Yeh *et al*., 2016), the cyclic nucleotide-gated ion channels CNGC2 and CNGC4 (Tian *et al*., 2019), as well as the plasma membrane calcium ATPases ACA8 and ACA10 (Frei dit Frey *et al*., 2012) (Table S1). These proteins were not unexpected as we had previously shown that RBOHD associates with EFR and FLS2 even before PAMP recognition (Kadota *et al*., 2014). Consistent with the observation that BAK1 interacts with EFR and FLS2 in a ligand-dependent manner (Chinchilla *et al*., 2007; Heese *et al*., 2007; Schulze *et al*., 2010; Roux *et al*., 2011; Sun *et al*., 2013), BAK1 was present in higher amounts after PAMP treatment (Fig. **S1b**; Table S1). These results show that the methods employed are effective for isolating active PRR-RBOHD complex(es) containing BAK1. We also identified proteins that are known to accumulate in the detergent-resistant plasma membrane fraction (DRM) after treatment with flg22, such as remorin and STOMATIN/PROHIBITIN/FLOTILLIN/HFLK/C (SPFH) domain containing proteins (Table S1)(Keinath *et al*., 2010). This result supports the observation in rice that RBOHs accumulate in the DRM after PAMP treatment (Nagano *et al*., 2016).

To identify proteins that play a role in the regulation of RBOHD, we selected 30 RBOHD-associated candidates based on their functions for transient expression assays to see what effect they would have on PAMP-induced ROS production. The candidate genes were expressed under the control of the CaMV 35S promoter in *N. benthamiana* and ROS production was measured after induction with flg22 (Fig. S2; Table S2). The effect of each candidate protein on flg22-induced ROS production was evaluated by comparing the regions where candidate proteins or GUS protein acting as a negative control were expressed in the same leaves under the control of the CaMV 35S promoter when introduced by Agroinfiltration. As positive control, we expressed BIK1. As expected, BIK1 expression resulted in a significant increase in flg22-induced ROS production (Fig. S2). Among 30 candidates, six genes significantly suppressed flg22-induced ROS production, and none increased it (Table S2). The six candidate negative regulators are KARYOPHERIN ENABLING THE TRANSPORT OF THE CYTOPLASMIC HYL1 (AT5G19820) (Zhang *et al*., 2017; Xiong *et al*., 2020), 2-oxoglutarate dehydrogenase E1 component (AT3G55410), lipase/lipoxygenase plat domain protein 2 (AT2G22170), an LRR-RK (AT3G02880), N-TERMINAL-TRANSMEMBRANE C2 DOMAIN PROTEINS TYPE 4/Ca^2+^-DEPENDENT LIPID-BINDING PROTEIN 1/SYNAPTOTAGMIN 7 (AT3G61050) (de Silva *et al*., 2011; Ishikawa *et al*., 2020; Lee *et al*., 2020b), and PB1CP (AT2G01190). PB1CP is the focus of the present study because a PB1 domain is conserved in p40phox and p67phox, which are important regulators of NOX2 in animals. The PB1 domain functions as a protein-binding module through PB1-mediated heterodimerization or homo-oligomerization. For example, the p40^phox^ and p67^phox^ proteins interact with each other through their PB1 domains, which facilitate the assembly of NOX2 and the cytosolic regulators p22^phox^, p40^phox^, and Rac at the membrane. Such assembly results in NOX2-mediated production of O_2_ ^-^ (Canton & Grinstein, 2014). Co-immunoprecipitation with FLAG-RBOHD indicated a total of three unique peptides corresponding to PB1CP from untreated, elf18, and elf18+flg22-treated samples (Fig. S1; Table 1). The expression of *PB1CP* under the control of the CaMV 35S or native promoters significantly reduced flg22-induced ROS production in *N. benthamiana* (Fig. 1, S3). These results suggest that PB1CP negatively regulates flg22-induced ROS production.

**Table 1.** Peptide counts of PB1CP in FLAG-RBOHD Co-IP analysis.

| PB1CP peptides identification by co-immunoprecipitation of FLAG-RBOHD and LC-MS/MS Analysis |  |  |  |  |
| --- | --- | --- | --- | --- |
| Treatment | Peptide Sequence | Probability | Best Mascot Score | Number of Spectra |
| mock | (K)SDDWFLNALNSAGLLNR(G) | 100% | 51.56 | 3 |
| elf18 | (K)SDDWFLNALNSAGLLNR(G) | 100% | 40.03 | 2 |
| elf18+flg22 | (K)SDDWFLNALNSAGLLNR(G) | 100% | 32.69 | 1 |
|  | (R)LLGLDDALALR(S) | 99% | 35.49 | 2 |
|  | (R)VHVEEPGGVR(T) | 96% | 20.71 | 1 |

**Fig. 1.**
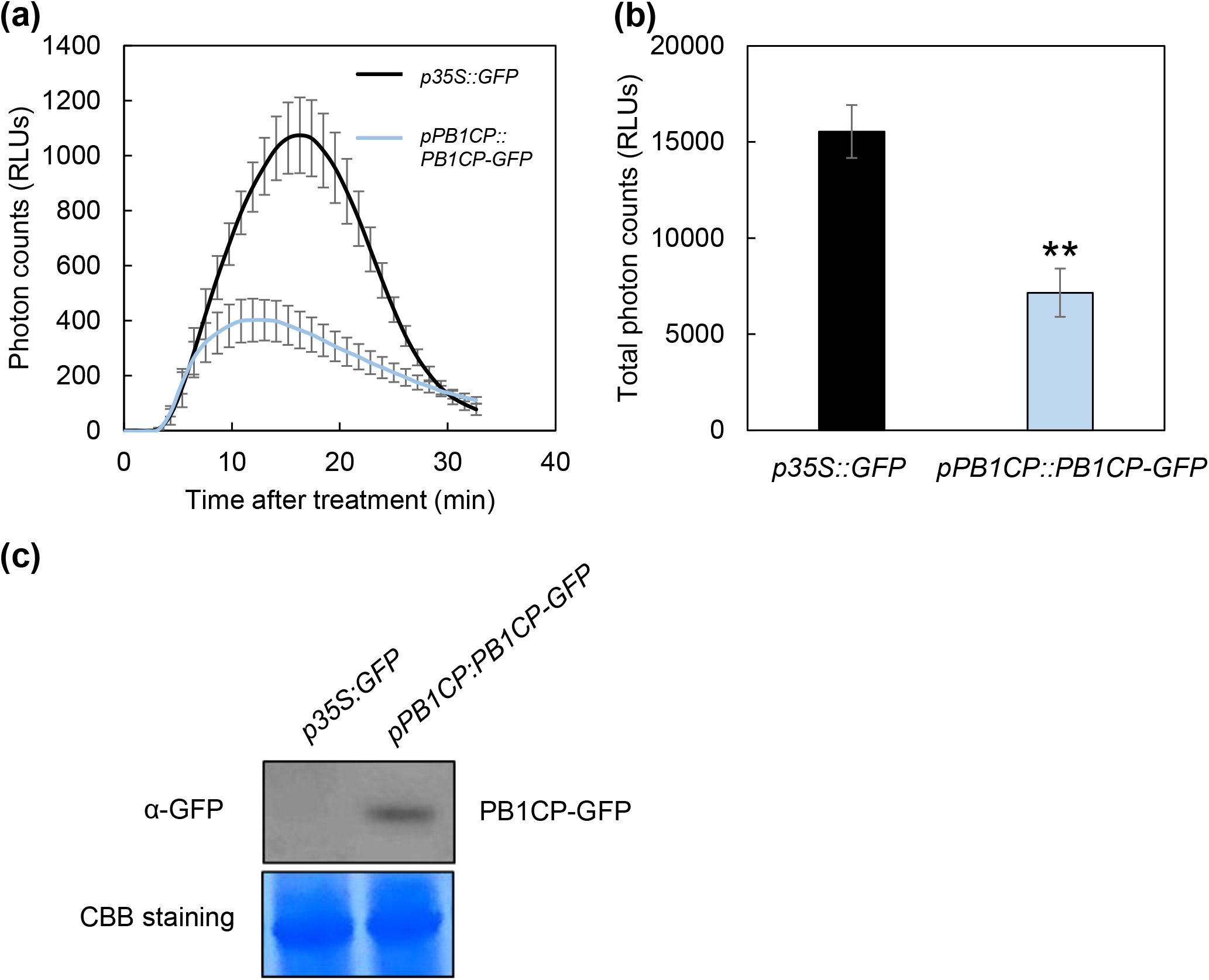
Heterologous expression of PB1CP-GFP under the native Arabidopsis promoter reduces flg22-induced ROS production in *N. benthamiana*. (**a, b**) The PB1CP-GFP (*pPB1CP::PB1CP-GFP*) and free GFP (*p35S::-GFP*) were expressed in the same leaf by Agroinfiltration, and flg22-induced ROS was measured in a luminol-based assay. Thirty minute time-course (**a**) and the total amount (**b**) of flg22-induced ROS production. Experiments were performed three times. Asterisks indicate a significant difference based on Student’s t-test (^∗∗^*p*-value ≤ 0.01). (**c**) The protein expression of PB1CP-GFP was confirmed by immunoblot analysis with α-GFP antibody (ab290; 1:8,000; Abcam).

### PB1CP negatively regulates PAMPs-induced ROS production in Arabidopsis

To clarify the role of PB1CP during PTI, we characterized two independent *pb1cp* mutants, *pb1cp-1* (SALK_036544) and *pb1cp-2* (SALK_207053). The *pb1cp-2* allele has a T-DNA insertion at the second exon, resulting in a potentially null mutant (Fig. **S4a**). In contrast, the *pb1cp-1* allele is unlikely null as it has T-DNA insertion within the 5’ UTR that results in a significant reduction in *PB1CP* transcript levels (Fig. **S4b**). These *pb1cp* mutants did not show any obvious phenotypic abnormalities (Fig. **S4c**). Importantly, the *pb1cp* mutants produced significantly higher ROS production upon induction with flg22, elf18, or chitin, than in Col-0 (Fig. **2a–c**). The *pb1cp* mutants did not show any difference in flg22-induced MAPK activation (Fig. **2d**), which is a ROS-independent signaling event during PTI (Shinya *et al*., 2014), suggesting that PB1CP is specifically involved in ROS production.

To further investigate the role of PB1CP during PTI, we generated two independent Arabidopsis transgenic lines overexpressing PB1CP-3×HA under the control of the CaMV 35S promoter (*p35S::PB1CP-3*×*HA*) (Fig. S5). Similar to the *pb1cp* mutants, neither *p35S::PB1CP-3*×*HA* line differed phenotypically from wild-type (Fig. **S5b**). Opposite to the *pb1cp* mutants, *p35S::PB1CP-3*×*HA* lines induced less ROS production upon treatment with flg22, elf18, or chitin, compared to wild-type Col-0 (Fig. **3a-c**), and flg22-induced MAPKs activation was unchanged in the transgenic lines (Fig. **3d**). Based on these results, we concluded that PB1CP specifically negatively regulates RBOHD-mediated ROS production in Arabidopsis.

### PB1CP reduces resistance against *C. higginsianum*

To test if there is a link between PB1CP-mediated regulation of ROS production and disease resistance, we measured resistance against the weakly virulent bacterial strain *Pseudomonas syringae* pv. *tomato* (*Pto*) DC3000 *COR*^-^ that lacks the toxin coronatine (COR) which otherwise triggers stomatal reopening during infection (Melotto *et al*., 2006), and the non-adapted bacterium *Pseudomonas syringae* pv. *cilantro* (*Pci*) 0788-9, which grows very poorly on Arabidopsis Col-0 plants (Lewis *et al*., 2008). Six-week-old Arabidopsis plants were spray-inoculated with *Pto* DC3000 *COR*^*-*^ and *Pci* 0788-9. There was no difference in bacterial growth of *Pto* DC3000 *COR*^-^ and *Pci* 0788-9 in the *pb1cp* mutants or *p35S::PB1CP-3*×*HA* lines, compared to Col-0 (Fig. **S6a,b**), suggesting that PB1CP is not required for resistance against these bacteria. Next, we quantified resistance against the fungal pathogen *C. higginsianum*. Six-week-old Arabidopsis leaves were drop-inoculated with *C. higginsianum*, and lesion diameters were measured (Fig. 4). Lesion diameters were smaller in the *pb1cp* mutants and larger in *p35S::PB1CP-3*×*HA* lines than in wild type, suggesting that *PB1CP* negatively regulates resistance against fungal infection, at least for a relatively host-specific pathogen like *C. higginsianum*.

**Fig. 2.**
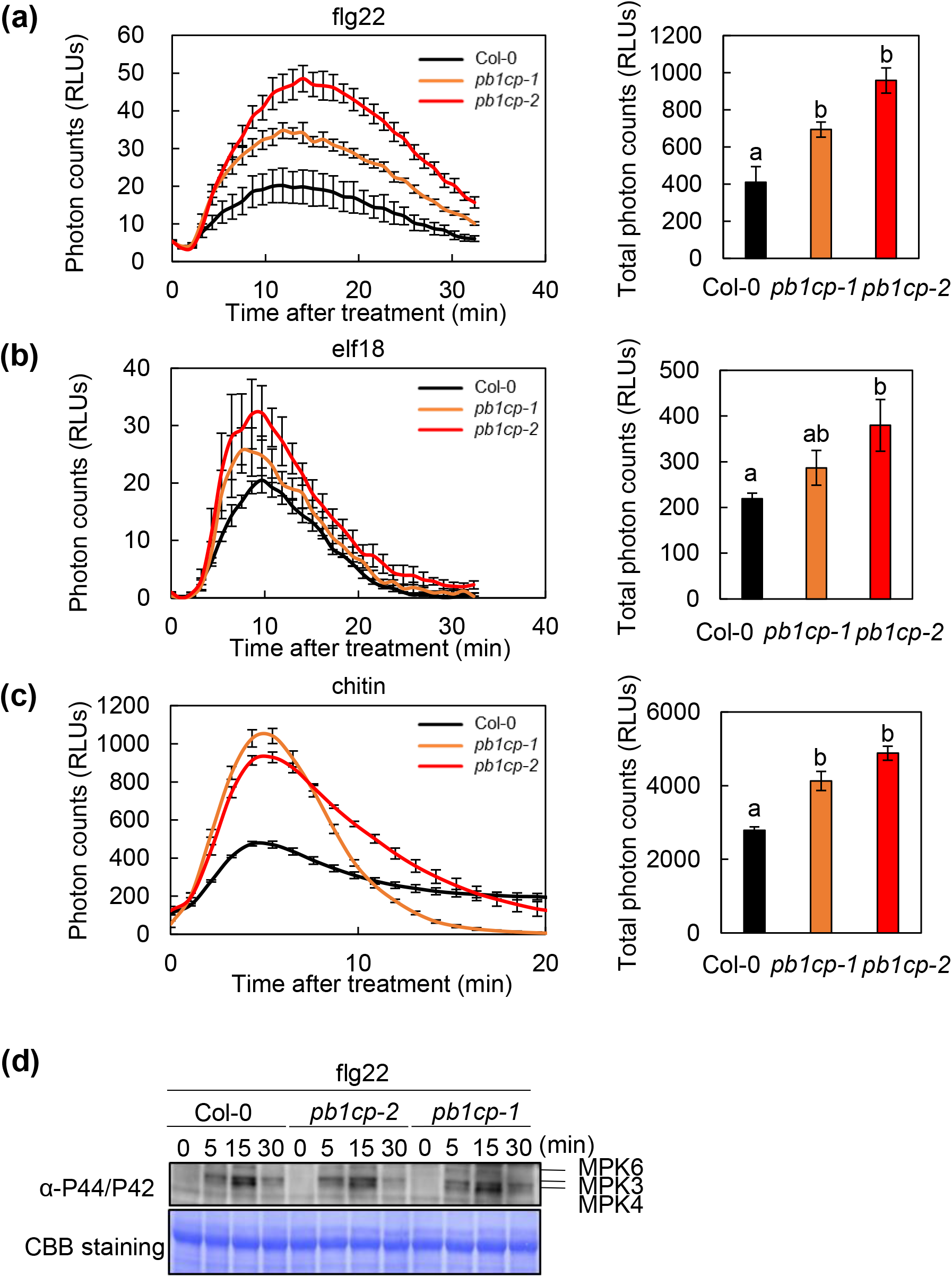
*pb1cp* mutants have higher PAMP-induced ROS production but normal MAPK activation Thirty-minute time-course and total amount of ROS production with flg22 (**a**), elf18 (**b**), or chitin (**c**) treatments of *pb1cp* mutants. The leaf discs of five-to six-week-old Arabidopsis were used for ROS assays. Different characters indicate significant differences based on one-way ANOVA and Tukey’s post hoc test (^∗^*p*-value ≤ 0.05). (**d**) flg22-induced activation of MAPKs in *pb1cp* mutants. Ten-day-old Arabidopsis seedlings were treated with flg22 and phosphorylated MAPKs were detected on immunoblots with α-phospho-p44/42 MAPK (Erk1/2) (Thr202/Tyr204) antibody (#4370; 1:2,000; Cell signaling Technology). Equal loading of protein samples is shown by CBB staining.

**Fig. 3.**
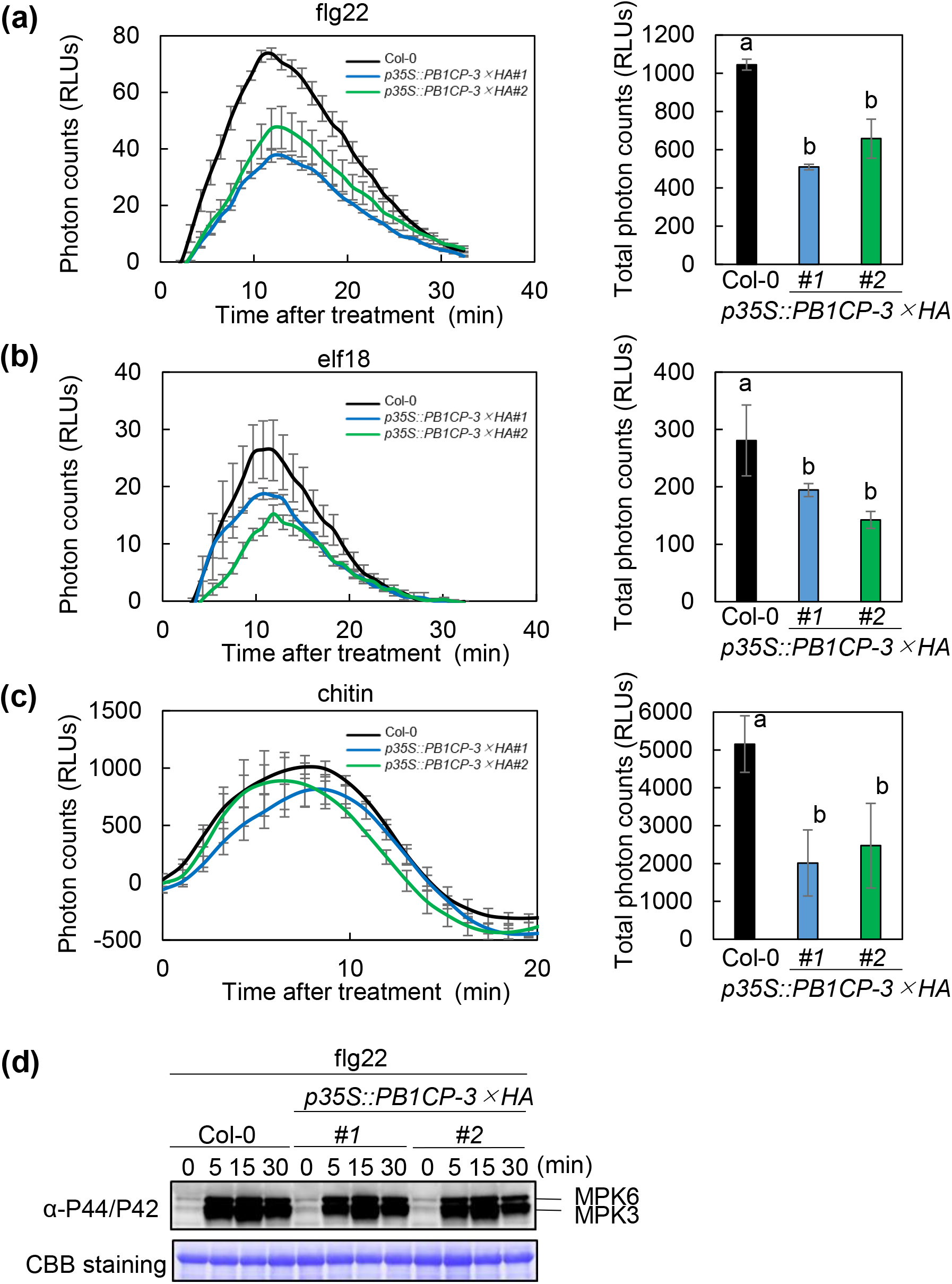
*PB1CP* overexpression lines have reduced PAMP-induced ROS production but normal MAPK activation. Thirty minute time-course and the total amount of ROS production with flg22 (**a**), elf18 (**b**), and chitin (**c**) treatment in *PB1CP* overexpression lines (*p35S::PB1CP-3*×*HA* #1 & #2). Leaf discs of five-to six-week-old Arabidopsis plants were used for ROS assays. Different characters indicate significant differences based on one-way ANOVA and Tukey’s post hoc test (^∗^*p*-value ≤ 0.05). (**d**) flg22-induced activation of MAPKs in *p35S::PB1CP-3*×*HA* lines. Ten-day-old Arabidopsis seedlings were treated with flg22, and phosphorylated MAPKs were detected on immunoblots with α-phospho-p44/42 MAPK (Erk1/2) (Thr202/Tyr204) antibody (#4370; 1:2,000; Cell signaling Technology). Equal loading of protein samples is shown by CBB staining.

**Fig. 4.**
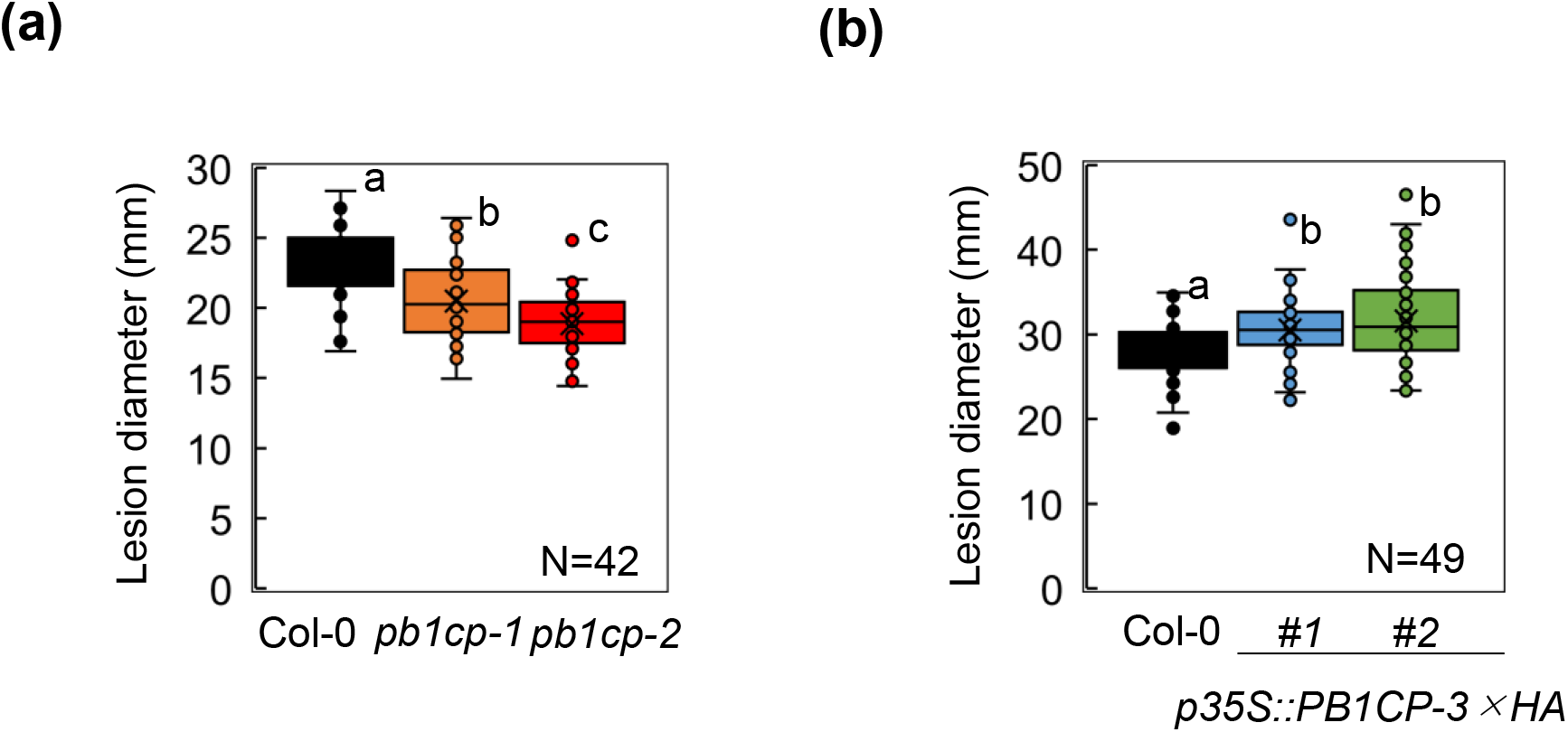
PB1CP suppresses resistance against *Colletotrichum higginsianum*. Diameters of necrotic lesions caused by *C. higginsianum* infection of *pb1cp* mutants (**a**) and in *PB1CP* overexpression lines (*p35S::PB1CP-3*×*HA #1* and *#2*) (**b**). Leaves of four-week-old soil-grown Arabidopsis plants were drop-inoculated with *C. higginsianum*.

### PB1CP directly interacts with RBOHD-ND, and PAMP treatment enhance the interaction

To confirm PB1CP-RBOHD interaction, we immunoprecipitated PB1CP-3×HA with α-HA antibody from the *PB1CP-3*×*HA* (*p35S::PB1CP-3*×*HA*) Arabidopsis line. PB1CP associated weakly with endogenous RBOHD (Fig. **5a**). Consistent with the original observation that PB1CP-derived peptides are present in greater amounts in PAMP-treated samples, treatment with elf18 or flg22 for 10 min increased PB1CP-RBOHD association. There was a similar but weaker effect of elf18, perhaps due to reduced EFR expression and elf18 recognition in Arabidopsis roots (Wu *et al*., 2016). In contrast, BIK1 was not detected in the PB1CP complex even after treatment with elf18 or flg22, suggesting that BIK1 may not be present in the PB1CP-RBOHD complex.

An *in vitro* binding assay using recombinant proteins was used to test if PB1CP and RBOHD bind directly (Fig. **5b**). Although we were unable to obtain full-length PB1CP recombinant protein, we were able to express domains of PB1CP (ND, PB1 domain (PB1D), and CD) in *E. coli. In vitro* pull-down assays showed that PB1CP-CD, but not PB1CP-ND nor PB1D, bind directly with RBOHD-ND. Furthermore, PB1CP-CD competed with BIK1 for RBOHD-ND binding (Fig. **5c**), suggesting that PB1CP-CD and BIK1 share an overlapping binding region of RBOHD-ND. The competition of PB1CP with BIK1 for binding with RBOHD is consistent with the finding that BIK1 was not part of the PB1CP-RBOHD complex *in vivo* (Fig. **5a**).

### PAMP treatment induces PB1CP accumulation in endomembrane compartments

To understand further how PB1CP regulates RBOHD, we monitored its subcellular localization by transiently expressing *PB1CP-GFP* in leaves of *N. benthamiana* by Agroinfiltration under the control of its own promoter (*pPB1CP::PB1CP-GFP*). Confocal microscopy showed that PB1CP-GFP signal was distributed in the cell periphery and cytoplasm (Fig. 6). We also observed that small PB1CP-GFP signal foci moved around the cell periphery and within the cytoplasm (Video S1). Interestingly, treatment with flg22or chitin for 3-6 h reduced PB1CP-GFP signals in the cell periphery and cytoplasm, and many PB1CP-GFP signal foci appeared in those regions (Fig. **6b**; Video S2). The PB1CP-GFP foci that appeared after treatment with flg22 clearly co-localized with FM4-64 dye, an endocytic tracer, showing that the foci localized at endomembrane compartments (Fig. **6c**). These results suggest that PB1CP moves from the cytoplasm and cell periphery to endomembrane compartments in response to PAMPs.

**Fig. 5.**
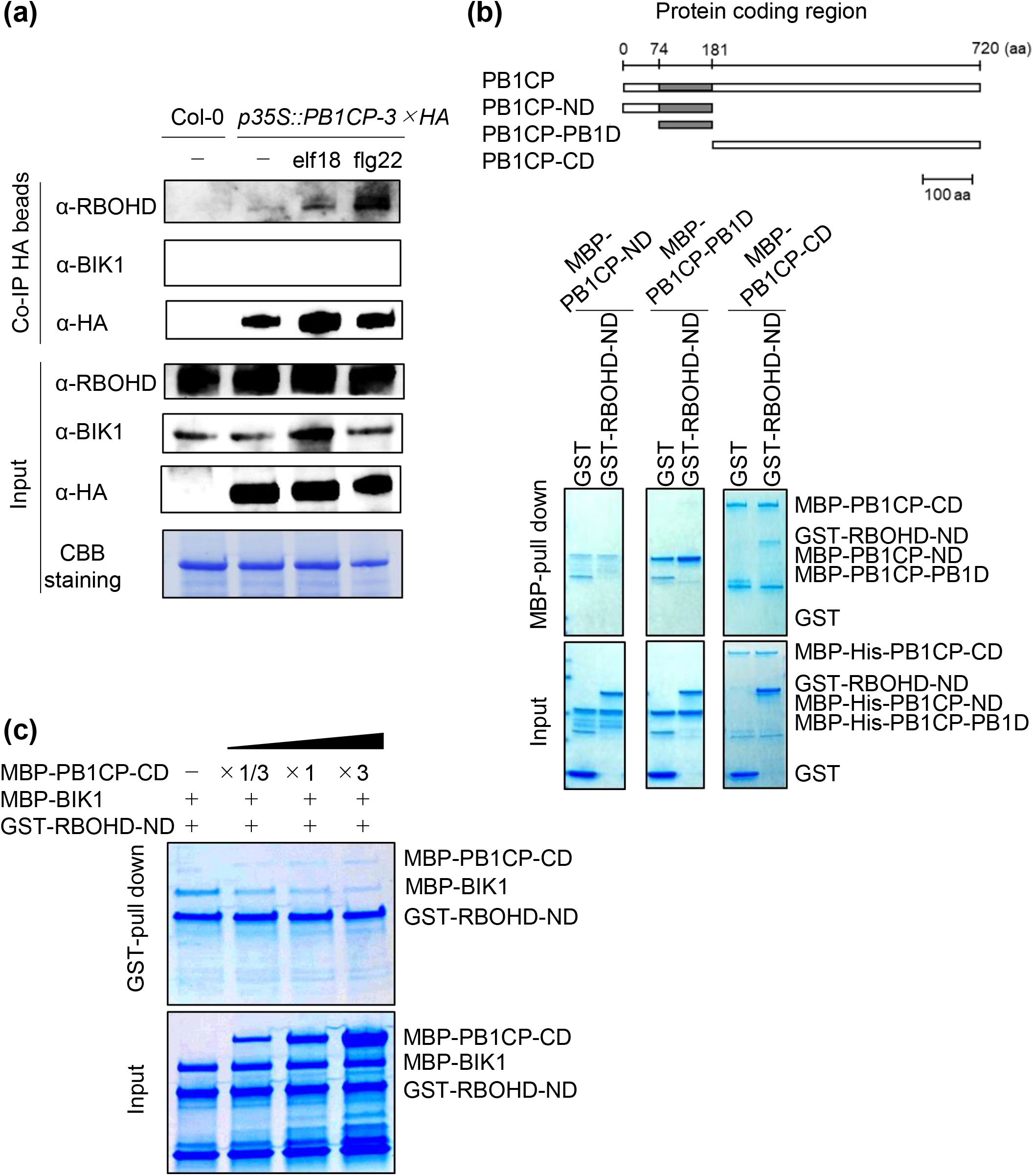
Treatment with elf18 or flg22 enhances PB1CP-RBOHD binding. (**a**) Co-immunoprecipitation of PB1CP and RBOHD in Arabidopsis. Stable transgenic Arabidopsis seedlings *p35S::PB1CP-3*×*HA* or Col-0 were treated with flg22 or elf18 for 10 min. Untreated plants (-) served as controls. Total proteins (input) were immunoprecipitated with α-HA magnetic beads followed by immunoblots with α-HA (3F10; 1:5,000, Roche), α-BIK1 (AS164030; 1:1,000; Agrisera), α-RBOHD (AS152962; 1:1,000; Agrisera) antibodies. Wild type Col-0 served as a negative control. (**b**) PB1CP-CD directly interacts with RBOHD-ND *in vitro*. MBP-PB1CP-ND, MBP-PB1CP-PB1D (PB1 domain), or MBP-PB1CP-CD were incubated with GST-RBOHD-ND or GST and with MBP. Input and pull-down proteins were separated by SDS-PAGE and stained with CBB. (**c**) PB1CP competes with BIK1 for binding to RBOHD. MBP-BIK1 was incubated with GST-RBOHD-ND with increasing amounts of MBP-PB1CP-CD, and pulled down with GST. All experiments were performed more than three times with similar results.

**Fig. 6.**
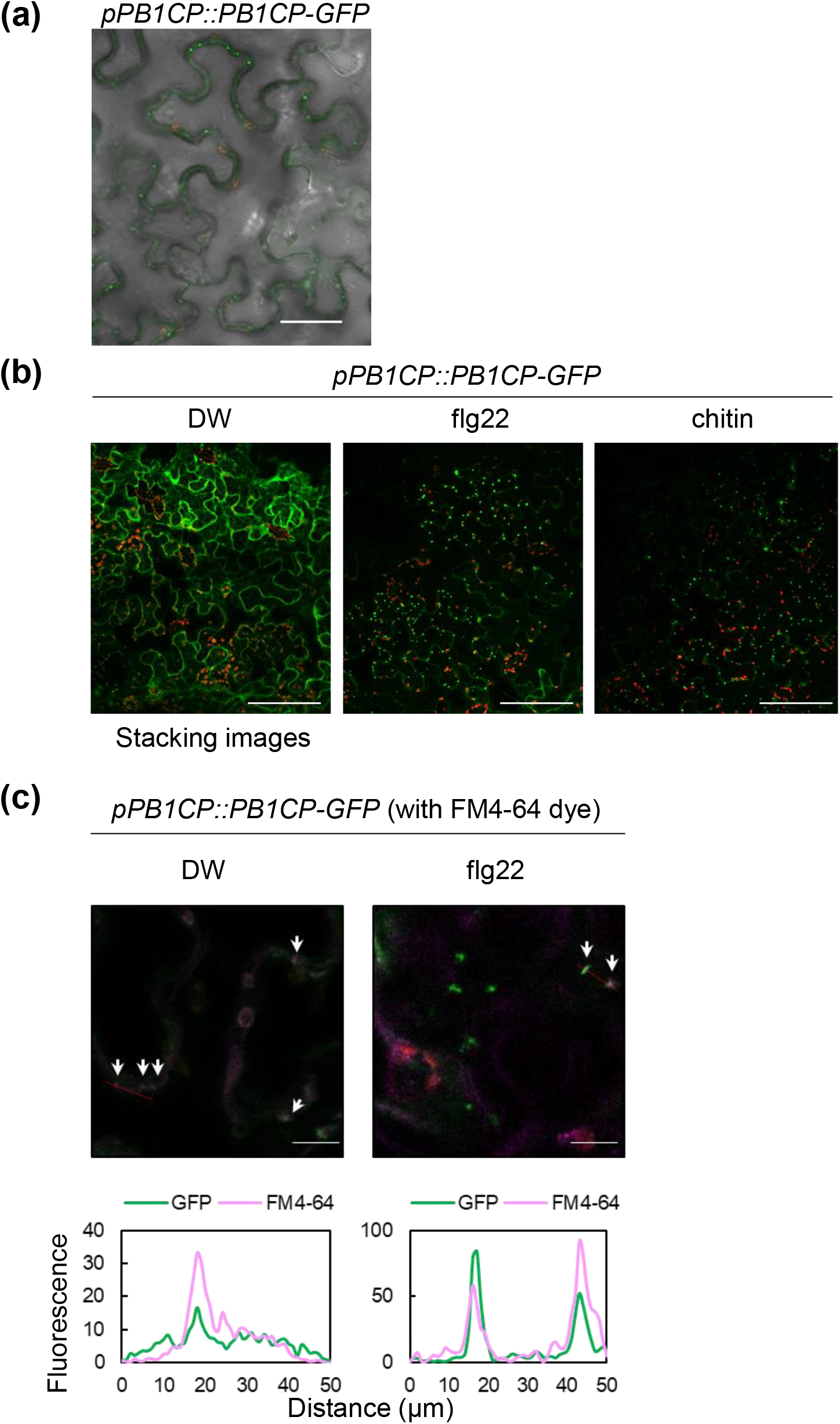
Subcellular localization of PB1CP in *N. benthamiana*. (**a**) Subcellular localization of PB1CP-GFP expressed under its native promoter in *N. benthamiana*. A white bar = 50 μm. (**b**) PAMPs (flg22 and chitin) induced the formation of PB1CP signal foci. PB1CP-GFP was expressed in *N. benthamiana* under the control of its native promoter. Leaf disks were treated with DW or 10 µM flg22 or 100 µM chitin for 3 h. White bars = 100 μm. (**c**) Co-localization of PB1CP-GFP and FM4-64 signals in cytoplasmic endomembrane compartments after treatment with flg22. Fluorescence intensities of PB1CP-GFP were quantified at 500 − 540 nm and FM4-64 at 558 −734 nm. Transections used for fluorescence intensity measurements are indicated by the red dash line. White bars = 30 μm.

**Fig. 7.**
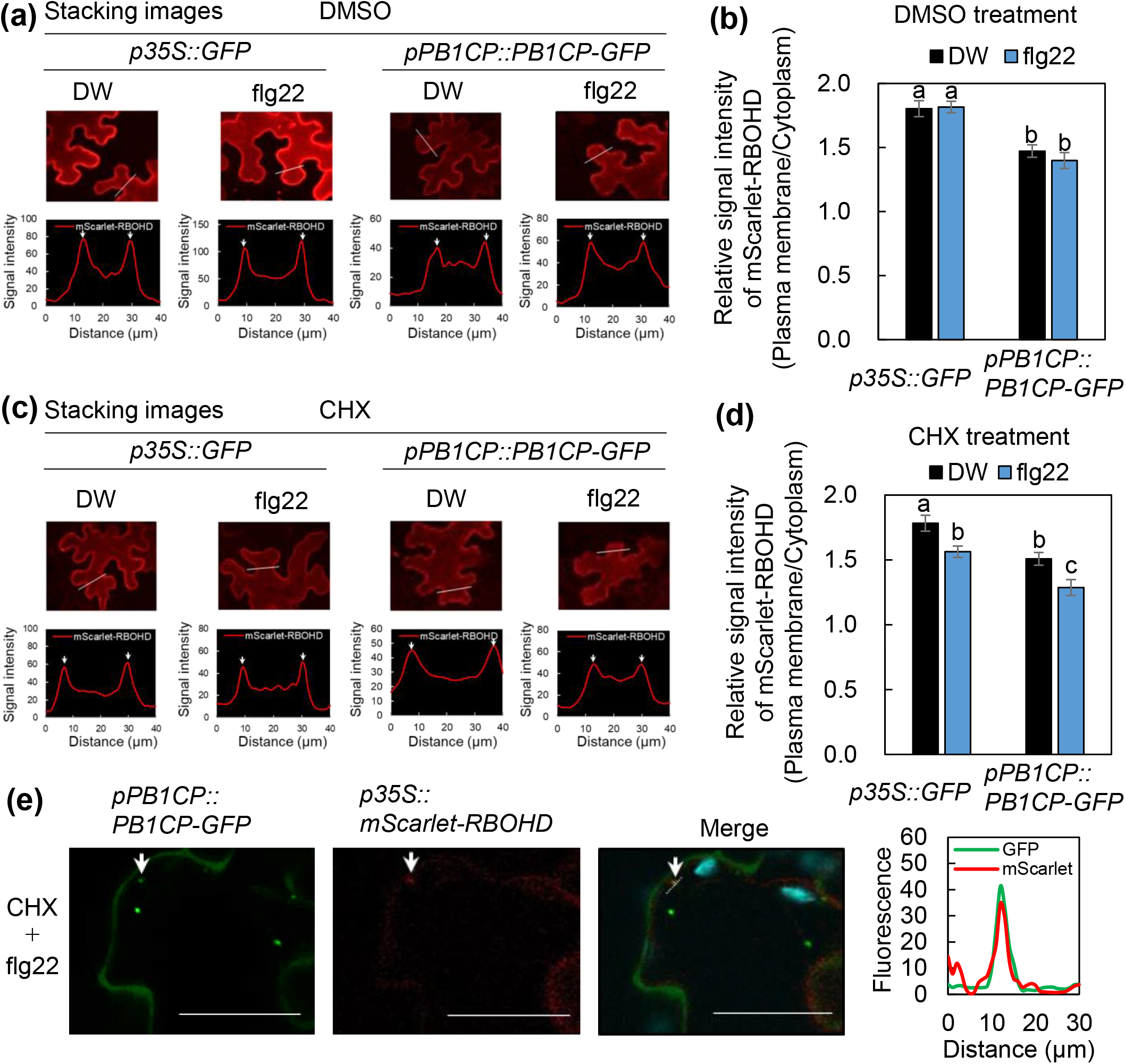
PB1CP reduces localization of RBOHD at the plasma membrane. (**a**) Localization of mScarlet-RBOHD (*p35S::mScarlet-RBOHD*) with co-expression of PB1CP-GFP (*pPB1CP::gPB1CP-GFP*) or GFP (*p35S::PB1CP-GFP*) in *N. benthamiana* after treatment with flg22. Distilled water served as treatment control. Fluorescence of mScarlet-RBOHD were measured at 559 − 595 nm. Transections used for fluorescence intensity measurements are indicated by broken white lines. (**b**) Intensity ratio of mScarlet-RBOHD at the plasma membrane to mScarlet-RBOHD at the cytoplasm. The data were extracted from ten cells from mScarlet-RBOHD image stacks. (**c**) Localization of mScarlet-RBOHD with co-expression of PB1CP-GFP or GFP after treatment with 300 μM CHX. Transections used for fluorescence intensity measurements are indicated by dashed white lines. (**d**) The intensity ratio of mScarlet-RBOHD at the plasma membrane to the cytoplasm after treatment with 300 μM CHX. The data was extracted from ten cells from mScarlet-RBOHD image stacks. Different letters indicate significantly different values at ^∗∗^*p* ≤ 0.05 (one-way ANOVA, Turkey’s *post hoc* test). Experiments were performed more than three times with similar results. (**e**) Co-localization of PB1CP-GFP and mScarlet-RBOHD signals in the endomembrane compartments in the cytoplasm after treatment with 300 μM CHX and 10 μM flg22. Fluorescence intensities of PB1CP-GFP were measured at 500 − 540 nm and mScarlet-RBOHD was measured at 559 − 595 nm. Transections used for fluorescence intensity measurements are indicated by the white dashed line. White bars = 100 μm.

### PB1CP reduces plasma membrane localization of RBOHD

We also determined RBOHD protein levels in *pb1cp* mutants and *p35S::PB1CP-3*×*HA* lines. Immunoblotting showed that the amounts of RBOHD were similar in all lines, suggesting that PB1CP does not affect RBOHD stability *in vivo* (Fig. **S8a**). RT-qPCR analysis also showed that transcript levels of *RBOHD* were similar (Fig. **S8b**). Since PB1CP-GFP accumulates within small endomembrane compartments (Fig. **6a,b**), we hypothesized that PB1CP regulates the endocytosis of RBOHD to control RBOHD protein levels at the plasma membrane. To test this hypothesis, we fused the mScarlet-tag-coding sequence to the 5’ end of the coding region of the genomic DNA of *RBOHD* (*gRBOHD*) and expressed mScarlet-RBOHD under the control of CaMV 35S promoter in *N. benthamiana*. This experiment confirmed that mScarlet-RBOHD is localized at the plasma membrane and is functional there (Fig. **S9a-c**). Importantly, we did not detect any truncation of mScarlet-RBOHD protein, confirming that the fluorescent signal is from full-length mScarlet-RBOHD (Fig. **S9d**). In a previous report, RBOHD expressed in tobacco cell cultures was localized at the plasma membrane as well as in intracellular compartments, mainly within Golgi cisternae (Noirot *et al*., 2014). There was no intracellular localization of mScarlet-RBOHD when expressed alone or co-expressed with free GFP in *N. benthamiana* (Fig. **7a, S9e**). However, co-expression with PB1CP-GFP reduced plasma membrane localization of mScarlet-RBOHD and increased its cytoplasmic localization (Fig. **7a,b**, **S9e**), suggesting a role for PB1CP in RBOHD relocalization from the plasma membrane to the cytoplasm.

Since PB1CP interaction with RBOHD is stronger after PAMP treatment, such treatment should further reduce the plasma membrane localization of RBOHD. However, there was no detectable relocalization of RBOHD upon flg22 treatment (Fig. **7a,b**). This may be due to newly synthesized RBOHD masking the effect of PB1CP. Thus, CHX was used to inhibit *de novo* synthesis of RBOHD. CHX treatment reduced the overall intensity of mScarlet-RBOHD, but mScarlet-RBOHD still clearly localized at the plasma membrane (Fig. **7c**). The expression of *PB1CP* reduced plasma membrane localization of RBOHD even in the presence of CHX. Importantly, flg22 treatment reduced plasma membrane localization in the presence of CHX (Fig. **7c,d**). Moreover, treatment with flg22 together with the expression of PB1CP-GFP further reduced plasma membrane localization of mScarlet-RBOHD in the presence of CHX (Fig. **7c,d**). These results suggest that PB1CP relocalizes plasma membrane-localized RBOHD upon PAMP treatment. It is also notable that mScarlet-RBOHD foci appeared in the cytoplasm of cells expressing PB1CP-GFP after treatment with CHX and flg22, and that some cytoplasmic foci co-localized with PB1CP-GFP (Fig. **7e**), indicating that there is a functional link between RBOHD and PB1CP *in vivo*.

## Discussion

### PB1CP negatively regulates RBOHD by direct binding

ROS produced by RBOHs serve as signaling molecules not only in plant immunity but also in a variety of biological processes such as abiotic stress responses, growth, and development (Suzuki *et al*., 2011). Plants tightly regulate the activity of RBOHs to minimize the detrimental effects of ROS, but the precise regulatory mechanisms of RBOH are unknown. In this work, we identified PB1CP as a previously unknown competitive binding protein for RBOHD. Phenotypic characterization of both *pb1cp* mutants and overexpressors showed that PB1CP negatively regulates RBOHD upon PAMP perception (Fig. **2a–c**, **3a–c**).

Because PB1CP negatively regulates RBOHD during PTI, we expected that loss- or gain-of-function lines of PB1CP would affect immunity against pathogens, but this was only true for the fungal pathogen *C. higginsianum* but not against the bacterial pathogens *Pto* DC3000 *COR*^*-*^ and *Pci* 0788-9 (Fig. 4, **S6a,b**). This may be because the amount of ROS required for defense against bacterial pathogens may be different than for fungal pathogens. For example, one of the primary defense mechanisms against these bacterial pathogens is based on stomatal closure (Melotto *et al*., 2008), which may not be affected when ROS levels are altered by PB1CP. In contrast, *C. higginsianum* has to pass through the plant cell wall and invaginate into the cell by modulating host plasma membrane during infection (O’Connell *et al*., 2012). It is possible that the fungus may be more sensitive to ROS because of its close association with the plasma membrane during this process. Alternatively, there may be some additional unknown fungal immunity mechanism controlled by ROS that is affected by PB1CP.

### PB1CP is in one of the eight groups of PB1 domain-containing proteins

Most of the PB1 domain ranges from 80 to 100 amino acids in length and exhibits a ubiquitin-like β-grasp fold with five β-sheets and two α-helices (Müller *et al*., 2006; Korasick *et al*., 2014). In animals, the PB1 domain functions as a protein-binding module for heterodimerization or homo-oligomerization. In particular, the PB1 domains of p40^phox^ and p67^phox^ interact with each other, which facilitates the assembly of other cytosolic regulators to NOX2 at the membrane (Groemping & Rittinger, 2005; Sumimoto, 2008). Arabidopsis encodes more than 80 PB1 domain-containing proteins, which can be segregated into eight families based on domain architecture (Mutte & Weijers, 2020). PB1CP belongs to the ‘kinase-derived family’, which is characterized by one PB1 domain in the ND with a large flanking sequence without any known domains. The PB1 domains of members in the ‘kinase-derived family’ resemble the PB1 domains of the ‘kinase domain family’. Kinase-derived PB1 domains appear to have been duplicated in the ancestors of angiosperms (Mutte & Weijers, 2020). The role of the PB1 domain in PB1CP still needs to be clarified, given that it is unlikely to be involved in interactions between RBOHD and PB1CP (Fig. **5b**). By analogy to p40^phox^ and p67^phox^, the PB1 domain of PB1CP may induce heterodimerization with an unidentified regulator of RBOHD. For instance, it is possible that the PB1 domain of PB1CP interacts with members of the ‘kinase-derived family’ or the ‘kinase domain family’. As evidence for this possibility, a yeast two-hybrid analysis showed that PB1CP binds to AT3G48240, another member of ‘kinase-derived PB1 family’, but the role of AT3G48240 in RBOHD regulation, if any, remains to be demonstrated (Arabidopsis Interactome Mapping Consortium 2011). Another possible role of the PB1 domain is to induce homo-oligomerization of PB1CP, which may result in the recruitment of RBOHD to the hypothetical PB1CP homo-oligomer.

Some PB1 domains can also undergo non-canonical interactions with proteins that do not have PB1 domains. For example, PAL OF QUIRKY (POQ), a PB1 domain-containing protein which belongs to the same group as PB1CP, interacts with STRUBBELIG (SUB), a cell surface LRR-RK, and QUIRKY (QKY), a protein containing multiple C2 domains and transmembrane regions (Trehin *et al*., 2013). Interestingly, POQ localizes at the cell periphery and in small cytoplasmic compartments, which is similar to PB1CP (Trehin *et al*., 2013). In addition, SUB is ubiquitinated *in vivo* and undergoes clathrin-mediated endocytosis (Gao *et al*., 2019). It is not known whether POQ is involved in the endocytosis of SUB, but it would be useful to compare the functions of PB1CP, POQ, and other PB1 domain-containing proteins in the same group during endocytosis.

### A model for the PB1CP regulatory mechanism of RBOHD

Although PB1CP associates with RBOHD, the interaction is more evident upon PAMP treatment (Fig. **5a, b**), suggesting that PB1CP binding is stronger with an activated form of RBOHD. Based on this result, we propose a model for PB1CP-mediated regulation of RBOHD in which PAMPs recognition triggers RLCK proteins such as BIK1 and CPKs to phosphorylate RBOHD-ND (Fig. S10) (Dubiella *et al*., 2013; Kadota *et al*., 2014; Li *et al*., 2014). At the same time, the EF-hand motifs at RBOHD-ND bind to Ca^2+^, whose entry into the cell is mediated by plasma membrane Ca^2+^ channels activated by BIK1 (Ogasawara *et al*., 2008; Oda *et al*., 2010). It is possible that PB1CP recognizes a phosphorylated form of RBOHD-ND or its structural changes caused by Ca^2+^ interaction. Alternatively, PB1CP may specifically bind to the homodimer of RBOHD at the plasma membrane once induced by PAMP treatment. In either case, once PB1CP interacts with RBOHD, BIK1-RBOHD binding is likely disrupted by both competition (Fig. **5c**) and the absence of BIK1 from the PB1CP-RBOHD complex (Fig. **5a**). Therefore, a possible function of PB1CP in the regulation of RBOHD is to release BIK1 from RBOHD, especially after activation by PAMPs. However, further examination of the activated RBOHD complex is required before this model can be confidently adopted.

Once PB1CP binds to RBOHD, it may induce relocation of the complex from the plasma membrane to the cytoplasm. This is suggested by the marked decrease in RBOHD levels at the plasma membrane and the increase at the cytoplasm following transient expression of PB1CP (Fig. **7a,b**). It is noteworthy that Agroinfiltration itself inevitably activates PTI via Agrobacterium-derived PAMPs. Thus it is possible that, at least in *N. benthamiana*, PB1CP binds tightly to RBOHD that is activated by PAMPs and relocates the protein. However, we cannot exclude the possibility that PB1CP controls plasma membrane RBOHD localization even in the absence of PAMPs when it is overexpressed.

The plasma membrane relocalization of RBOHD upon PAMP treatment was only detectable in the presence of CHX (Fig. **7c,d**) because mScarlet-RBOHD is highly expressed, and the supply of newly synthesized RBOHD to the plasma membrane would make it difficult to observe RBOHD dynamics. We tried to express mScarlet-RBOHD under the control of native promoter in *N. benthamiana*, but failed to detect fluorescence, probably because of low expression. In the presence of CHX, we detected RBOHD signal foci co-localized with PB1CP after PAMP treatment (Fig. **7e**), suggesting that RBOHD moves from the plasma membrane to the cytoplasm in concert with PB1CP. Clathrin- and microdomain-dependent endocytic pathways were shown to cooperatively regulate RBOHD dynamics (Hao *et al*., 2014). Thus, one attractive hypothesis is that PB1CP-mediated relocalization of RBOHD is through endocytosis, and the small mScarlet-RBOHD and PB1CP-GFP signal foci may be endosomes. We also detected the clathrin-related components HAP13 and AP4M, and flg22-inducible DRM components such as remorin and SPFH proteins by co-immunoprecipitation with RBOHD (Table S1). In addition, the E3 ubiquitin ligase PIRE, which interacts with both PBL13 and RBOHD, ubiquitinates RBOHD and decreases its abundance, possibly through endocytosis of RBOHD and vacuolar degradation (Lin *et al*., 2015; Lee *et al*., 2020a). Thus it is worth investigating the molecular relationship between PB1CP and PIRE for the endocytosis of RBOHD. It would also be interesting to test whether PB1CP-based regulation extends to other plant RBOHs, which have diverse functions in stress adaptation, growth, and development.

We have focused on the role of PB1CP in this work, but other candidate RBOHD-associated proteins are also of interest and will be explored elsewhere.

## Acknowledgments

We thank all members of the Shirasu lab for their discussion of and insights about the work. We thank Ms. Akiko Ueno, Ms. Naomi Watanabe, Ms. Mamiko Kouzai, Ms. Yoko Nagai, and Ms. Kanako Hori for their support of this project. We thank Dr. Max Fishman for critical reading of the manuscript. The research was financially supported by JSPS KAKENHI Grant Numbers 16J00771 (to Y.G.), 16H06186, 16KT0037, 20H02994 (to Y.K), 15H05959, 17H06172 (to K.S), as well as the Gatsby Charitable Foundation (F.L.H.M and C.Z.) and the European Research Council (project ‘PHOSPHinnATE’, grant agreement No. 309858).

## Author contributions

Y.G, Y.K, C.Z, and K.S. supervised the research. Y.K. performed co-immunoprecipitation of RBOHD. J.S, P.D, and F.L.H.M. performed LC-MS/MS analyses, N.M. helped generate constructs of RBOHD-associated proteins for transient expression in *N. benthamiana*, and Y.G. performed the other experiments. Y.G, Y.K, and K.S. wrote the manuscript. All of the authors, read, commented on, and approved the manuscript.

---

**Fig. S1.**
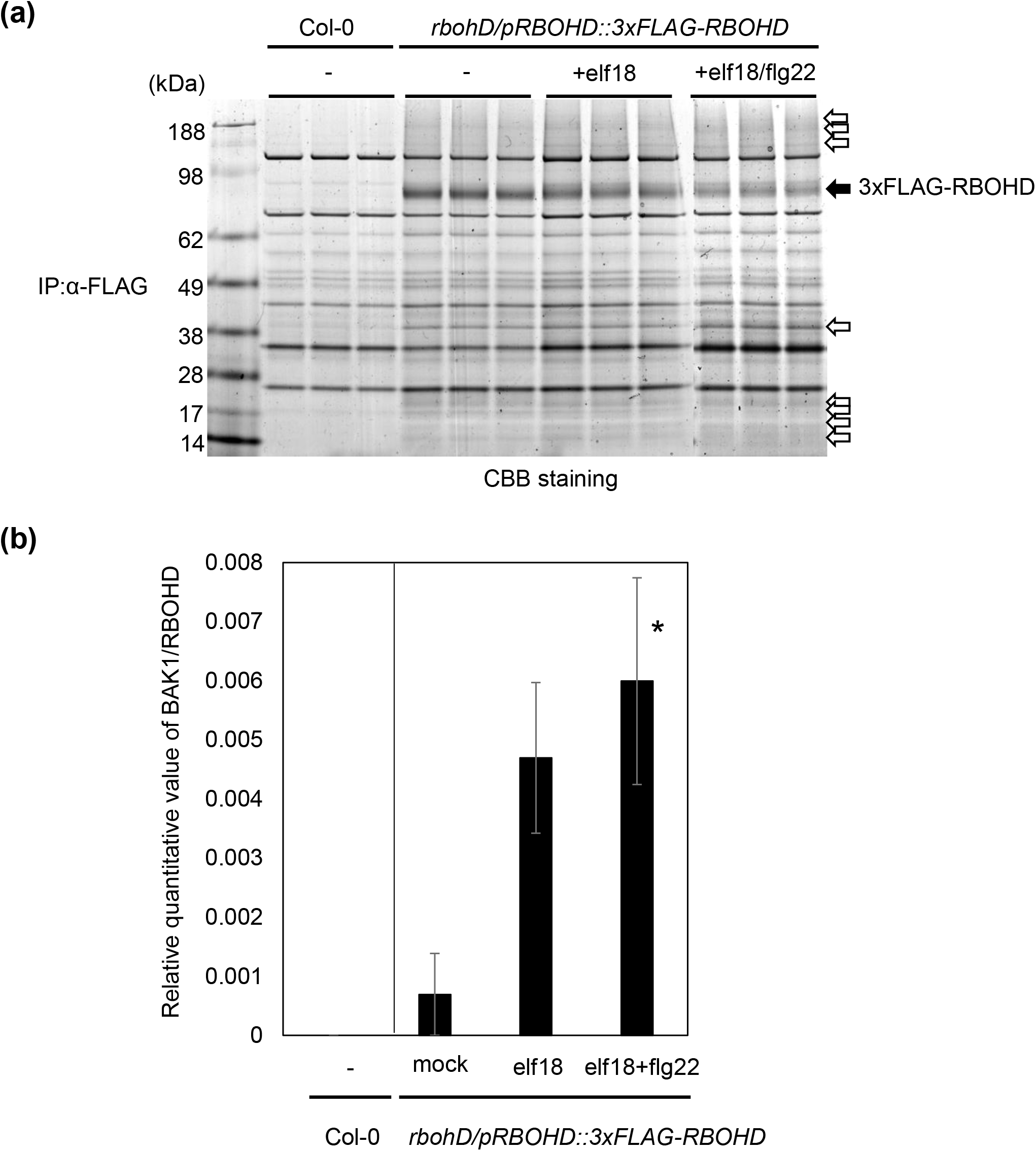
Co-immunoprecipitation of 3×FLAG-RBOHD to identify RBOHD-associated proteins. (**a**) SDS-PAGE gel of proteins enriched by α-FLAG immunoprecipitation stained with CBB. Co-immunoprecipitation with α-FLAG beads was performed using stable transgenic Arabidopsis plants expressing 3×FLAG-RBOHD under the control of its native promoter (*rbohD/pRBOHD::3*×*FLAG-RBOHD*) after treatment with DW, 1 µM elf18, or 1 µM elf18 and 1 µM flg22 simultaneously for 10 min. Non-transformed wild type Col-0 was used as a negative control. Black arrow indicates the 3×FLAG-RBOHD band. White arrows indicate proteins that specifically eluted with 3×FLAG-RBOHD. (**b**) Treatment with elf18 or elf18+flg22 increased the binding of BAK1 with RBOHD. BAK1 values were normalized against RBOHD. Data are means ± SE of three biological replicates. The asterisk indicates significant differences among mock, elf18 and elf18+flg22 samples at ^∗^*p* ≤ 0.05 (one-way ANOVA, Dunnett’s post hoc test).

**Fig. S2.**
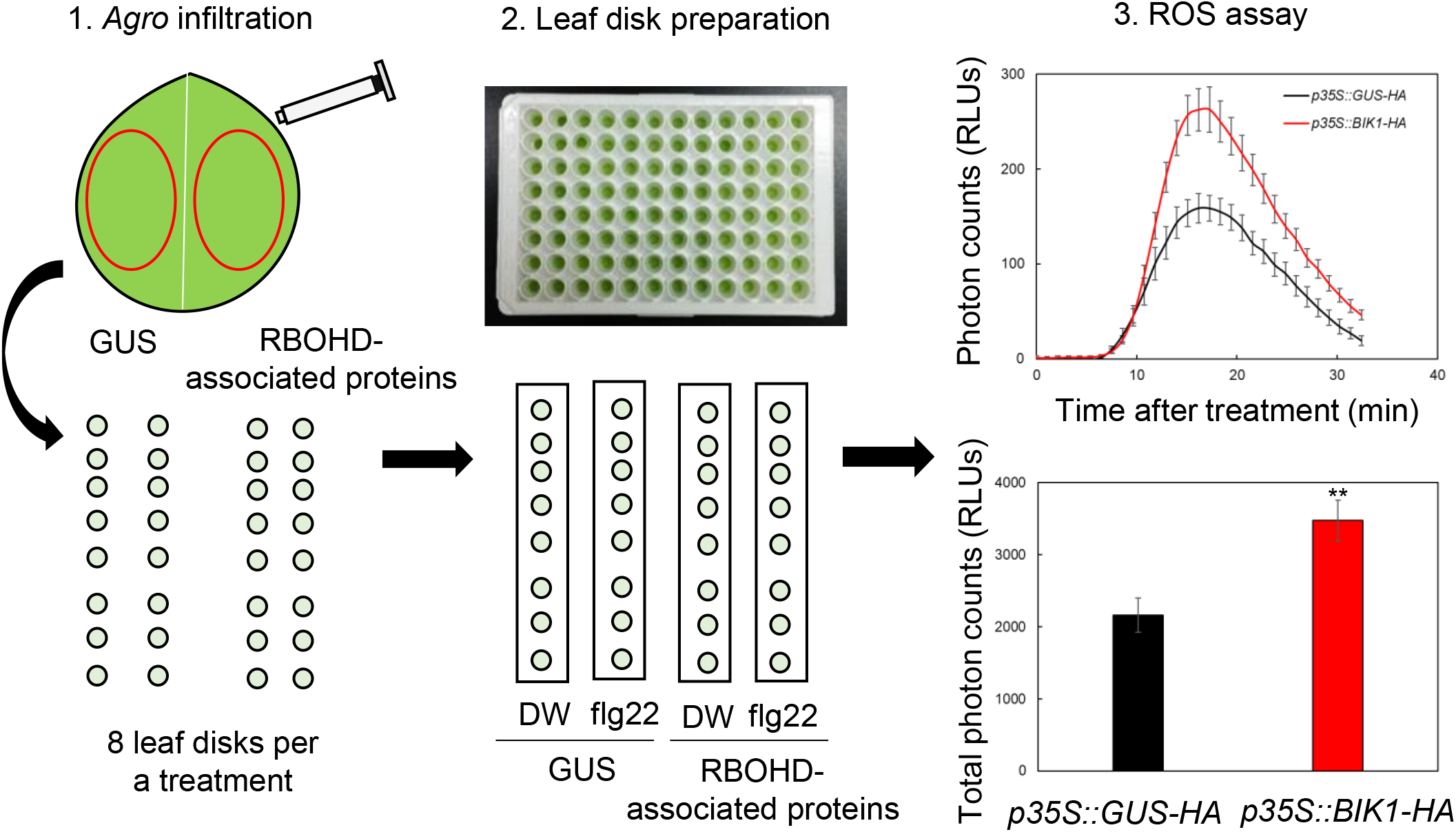
Rapid functional analyses of RBOHD-associated proteins in *N. benthamiana*. RBOHD-associated proteins fused with 3×HA at the C-terminus or BIK1-HA (positive control) and GUS-HA (negative control) were expressed under the control of the CaMV 35S promoter in the same leaf of *N. benthamiana* by Agroinfiltration. 30 min time-course and total amount of flg22-induced ROS were measured by luminol-based assays. Experiments were performed four times with similar results.

**Fig. S3.**
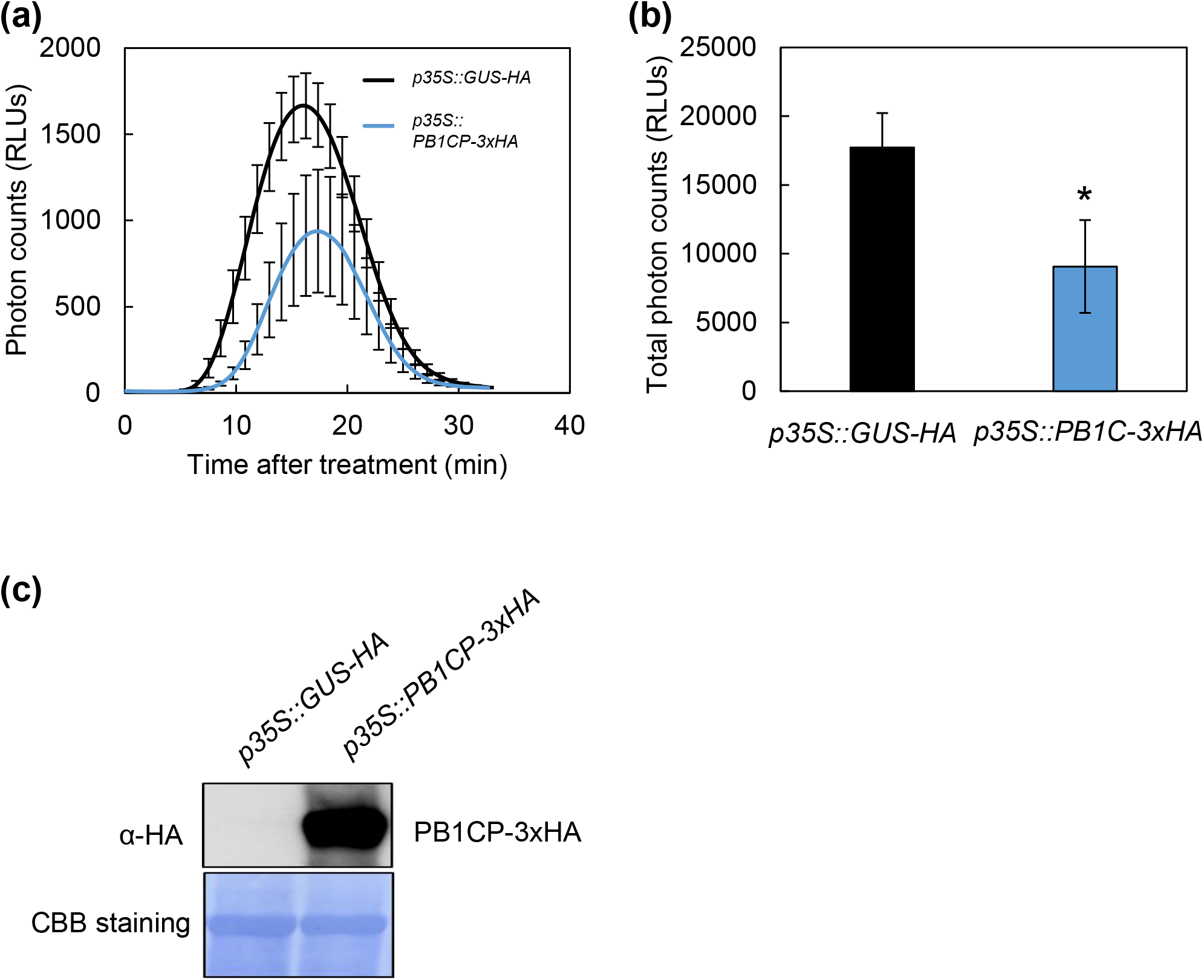
PB1CP inhibits flg22-induced ROS production in *N. benthamiana. PB1CP-3*×*HA* and *GUS-HA* were expressed under the CaMV 35S promoter in the same leaf by Agroinfiltration, and flg22-induced ROS was measured in a luminol-based assay. Thirty-minute time-course (**a**) and the total amount (**b**) of flg22-induced ROS production. Experiments were performed three times with similar results. Asterisks indicate significant differences based on Student’s t-test (^∗^*p*-value ≤ 0.05). (**c**) Protein expression of PB1CP-3xHA was confirmed by immunoblots with α-HA antibody (3F10; 1:5,000; Roche).

**Fig. S4.**
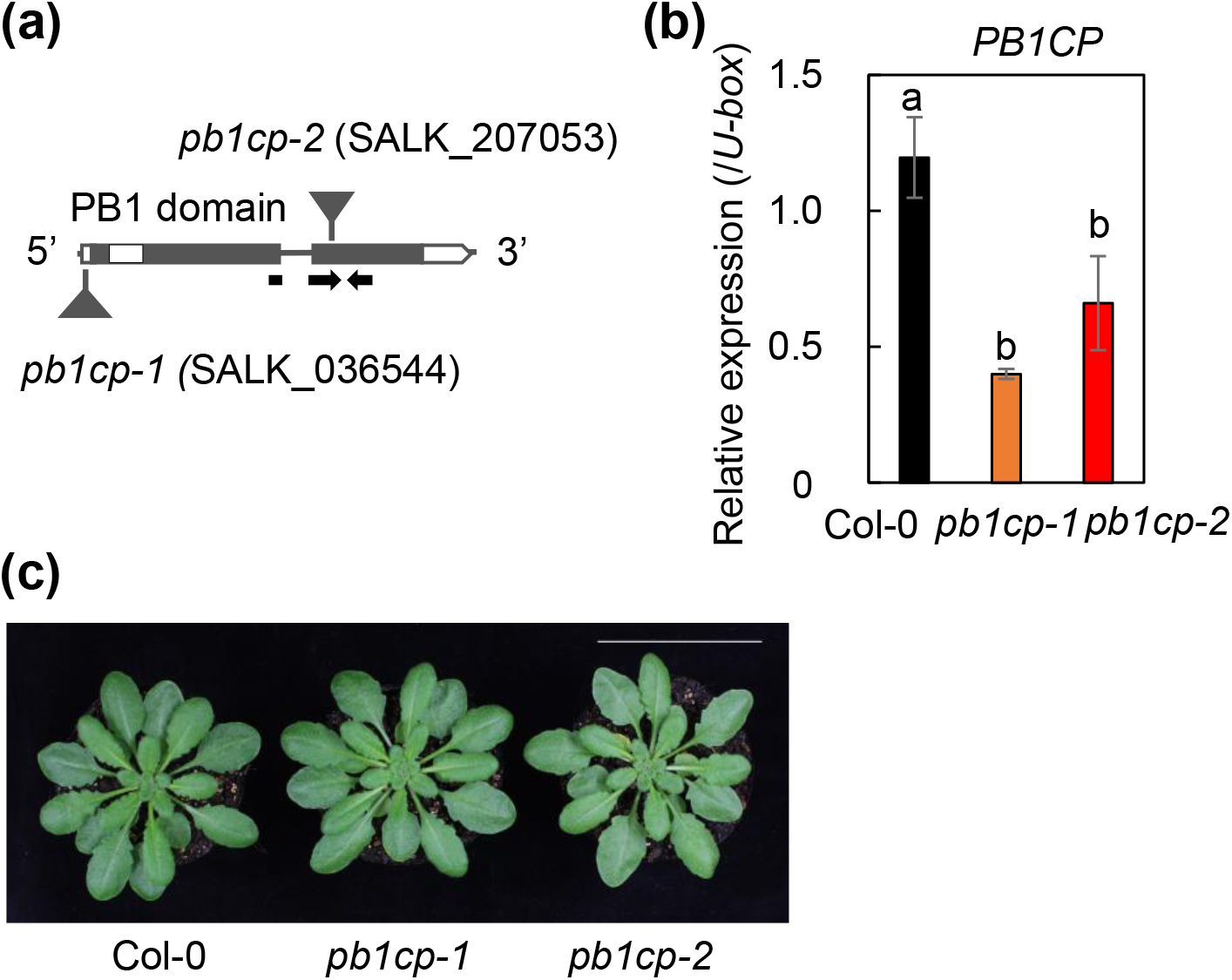
T-DNA insertion and expression in *pb1cp* mutants. (**a**) Positions of T-DNA insertions within the *PB1CP* locus in *pb1cp-1* (SALK_036544) and *pb1cp-2* (SALK_207053) alleles. (**b**) Transcript levels of *PB1CP* in the *pb1cp* mutants were measured by Quantitative RT-PCR (RT-qPCR) after normalization to the *U*-*box* housekeeping gene transcript (*At5g15400*). Different letters indicate significant differences at ^∗∗^*p* ≤ 0.01 (one-way ANOVA, Turkey’s *post hoc* test). (**c**) Phenotype of 6 week-old wild type and *pb1cp* mutants. A white bar = 5 cm.

**Fig. S5.**
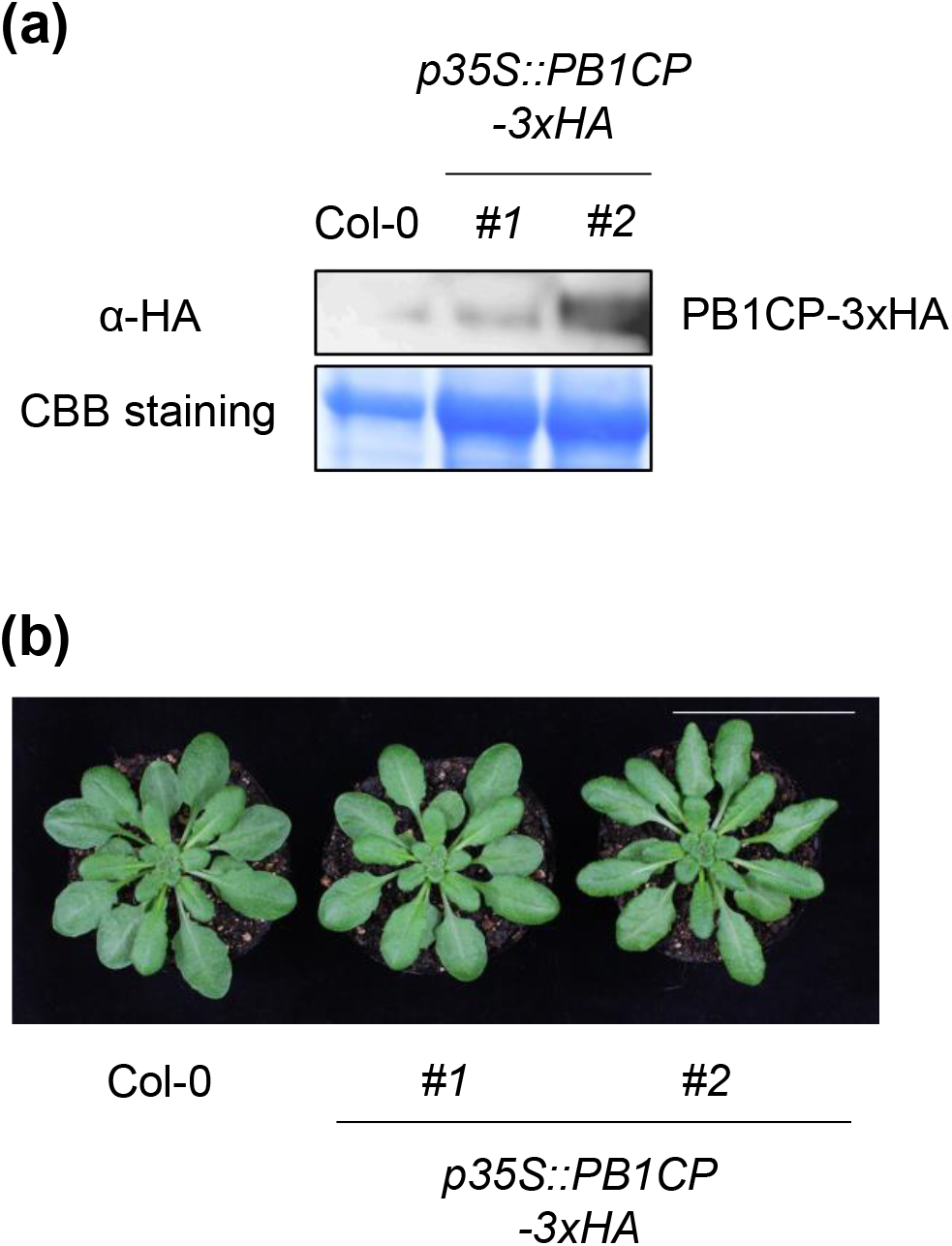
Expression in *PB1CP* overexpression lines. (**a**) PB1CP-3×HA protein in *p35S::PB1CP-3*×*HA* lines. (**b**) Phenotype of wild type and *p35S::PB1CP-3*×*HA* lines. Fourteen-day-old Arabidopsis seedlings were used to measure PB1CP-3×HA protein expression. PB1CP protein was detected with α-HA antibody (3F10; 1:5,000; Roche). Protein loading was visualized by CBB staining. A white bar = 5 cm.

**Fig. S6.**
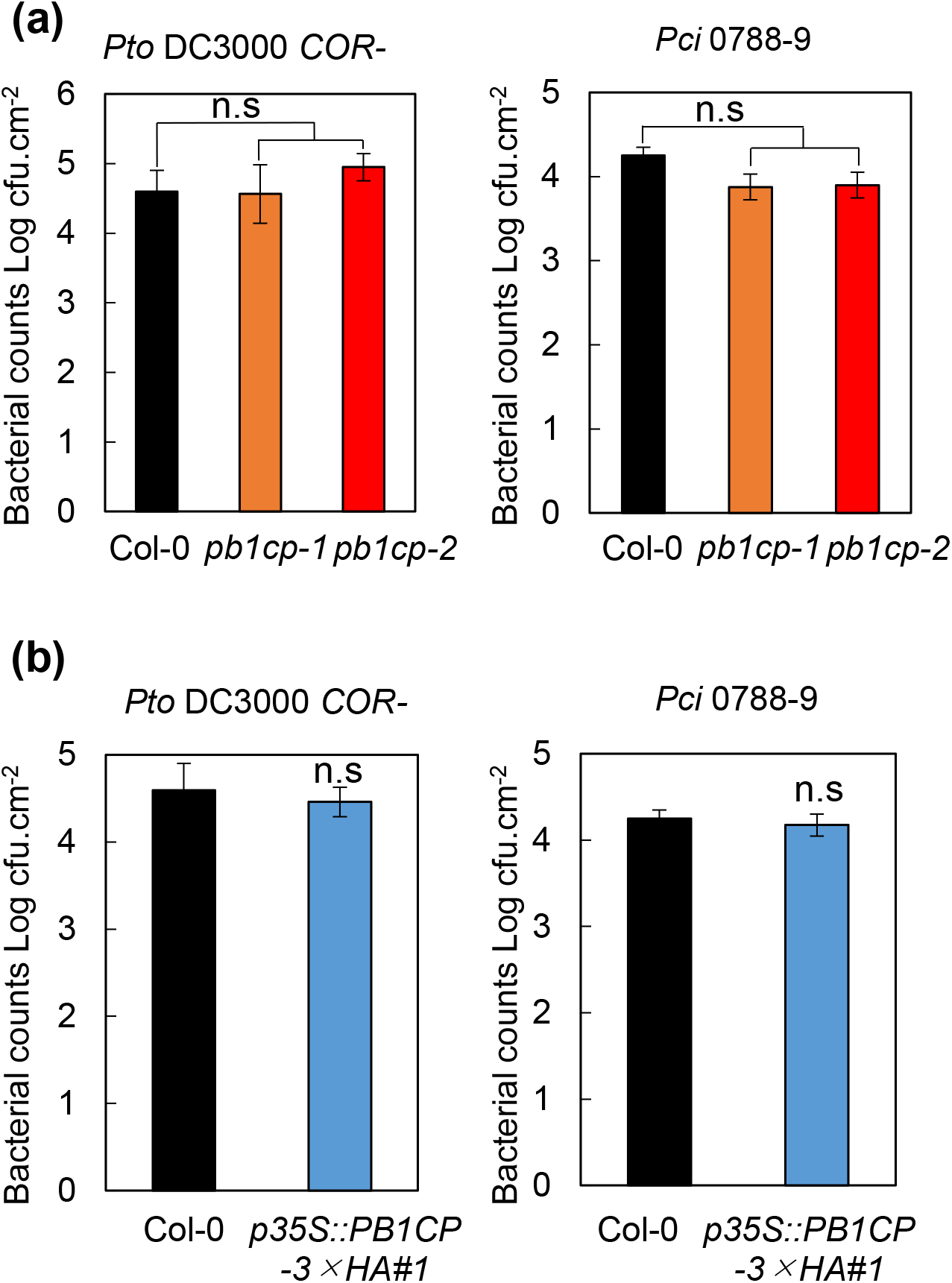
*pb1cp* mutants and *PB1CP* overexpression lines do not show defects in resistance against bacteria. *Pseudomonas syringae* pv. *tomato* DC3000 (*Pto* DC3000) lacking the toxin coronatine (*COR*-) (OD600 = 0.02) or *Pseudomonas syringae* pv. *cilantro* (*Pci*) 0788-9 (OD600 = 0.02) were sprayed onto leaf surfaces of *pbl1cp* mutants (**a**) and *p35S::PB1CP-3*×*HA* lines (**b**), and plants were maintained uncovered. Bacterial numbers (cfu) were determined 3 days post-inoculation. Values are means ± SE of six biological replicates.

**Fig. S7.**
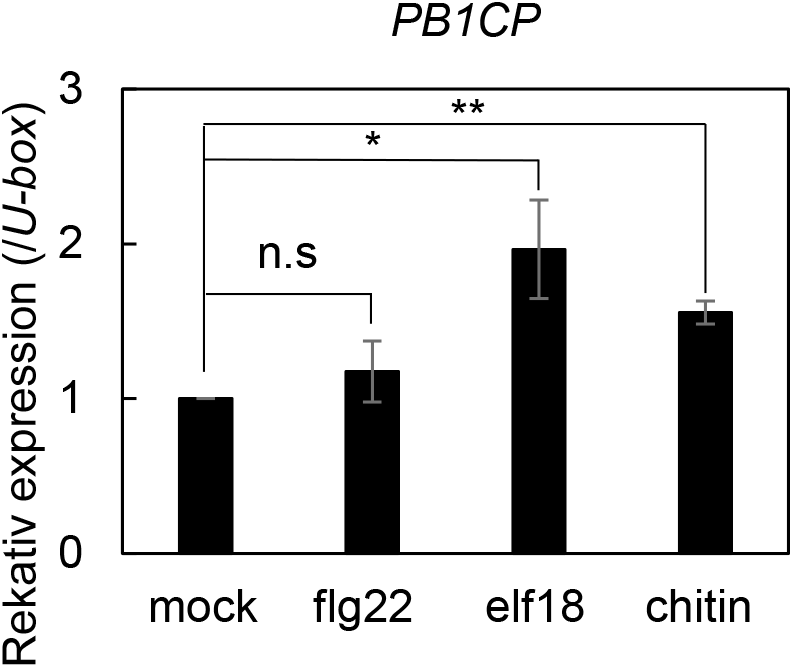
elf18 and chitin, but not flg22 weakly induce the accumulation of *PB1CP*. Transcript levels of *PB1CP* in Arabidopsis seedlings after treatment with DW (mock), 1 μM flg22, 1 μM elf18, or 10 μM chitin were measured by RT-qPCR after normalization to *PUB* housekeeping gene transcription (*At5g15400*). Data are means ± SE of three biological replicates. Asterisks indicate significant differences from mock treatment (Student’s t-test, ^∗^*p* ≤ 0.05; ^∗∗^*p* ≤ 0.01).

**Fig. S8.**
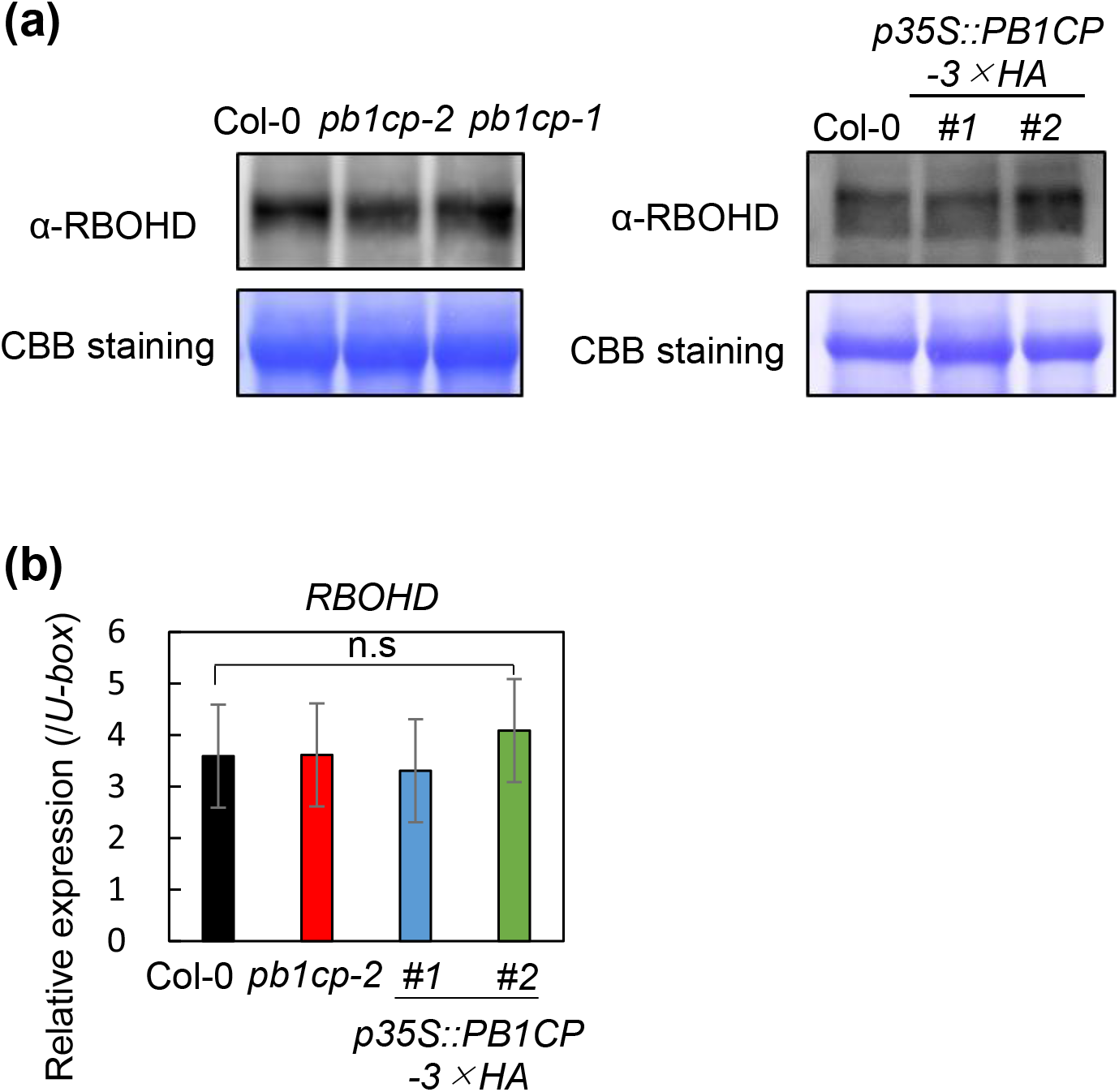
PB1CP does not affect expression of RBOHD. (**a**) Immunoblots of RBOHD protein in *pb1cp* mutants and *p35S::PB1CP-3*×*HA* lines. RBOHD protein was detected with α-RBOHD antibody (AS152962; 1:1,000; Agrisera). The equal loading of protein samples was shown by CBB staining. (**b**) Transcript levels of *RBOHD* in *pb1cp* mutants and *p35S::PB1CP-3*×*HA* lines were measured by RT-qPCR after normalization to the *PUB* housekeeping gene (*At5g15400*). Data are means ± SE of three biological replicates.

**Fig. S9.**
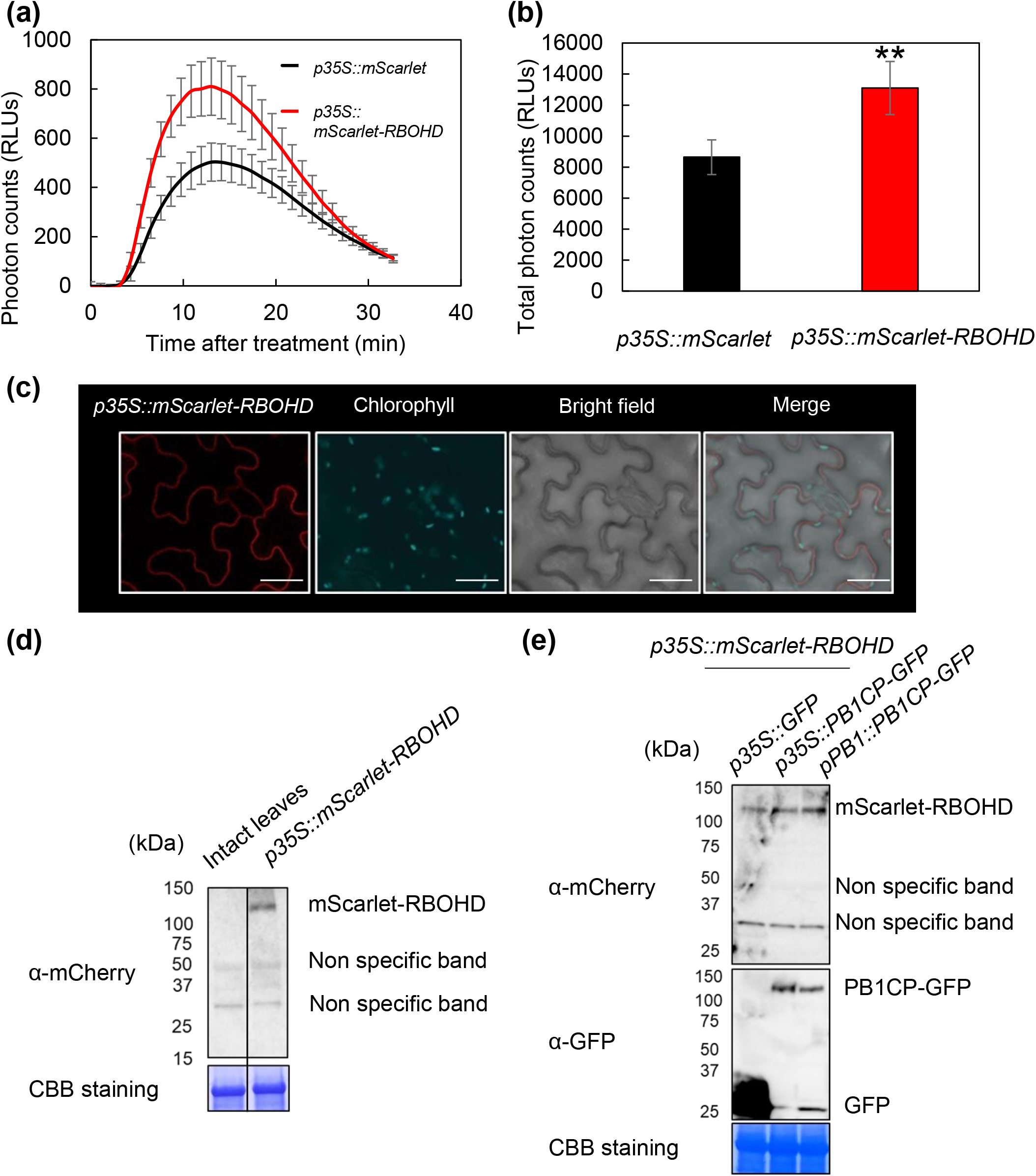
mScarlet-RBOHD is functional and is localized at the plasma membrane. (**a, b**) The overexpression of *mScarlet-RBOHD* increased flg22-induced ROS production in *N. benthamiana. mScarlet-RBOHD* and *mScarlet* were expressed under the CaMV 35S promoter in the same leaf by Agroinfiltration. ROS production was measured by luminol-based assay. Thirty minute time-course (**a**) and the total amount (**b**) of flg22-induced ROS production. The asterisk indicates significant differences at ^∗∗^*p* ≤ 0.01 (Student’s t-test). Experiments were performed three times with similar results. (**c**) Localization of mScarlet-RBOHD expressed under control of the CaMV 35S promoter (*p35S::mScarlet-RBOHD*) and chlorophyll autofluorescence. White bars = 30 μm. (**d, e**) The protein expression of mScarlet-RBOHD (*p35S::mScarlet-RBOHD*) and PB1CP-GFP (*p35S::PB1CP-GFP, pPB1CP::PB1CP-GFP*) in *N. benthamiana* were confirmed by immunoblots with α-mCherry (ab167453; 1:1,000; Abcam) and α-GFP (ab290; 1:8,000; Abcam) antibodies.

**Fig. S10.**
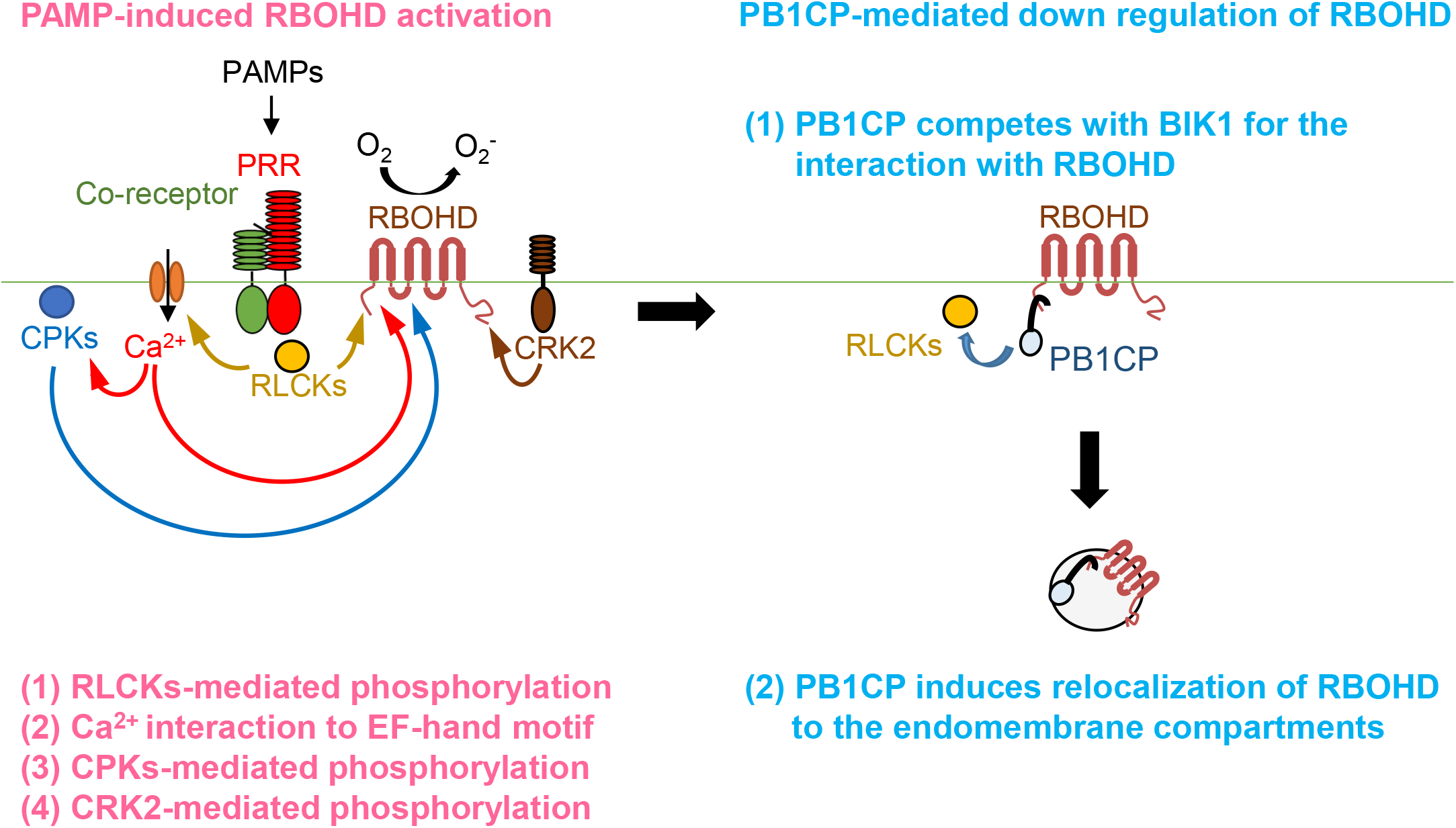
A model for PAMP-induced RBOHD activation and PB1CP-mediated RBOHD down-regulation during PTI. The left figure shows the RBOHD activation model during PTI. Upon PAMP perception, PRRs and co-receptors directly phosphorylate and activate RLCKs. The phosphorylated RLCKs bind the N-terminal domain of RBOHD and phosphorylate it on several specific sites (1). The RLCKs-mediated phosphorylation would mediate the Ca^2+^-based regulation of RBOHD by inducing conformational changes that could lead to increased Ca^2+^ binding affinity for EF-hand motifs and/or increased accessibility for CPK-mediated phosphorylation (2). At the same time, motif in RBOHD and also activation of CPKs, which phosphorylates the N-terminal domain of RBOHD (3). The produced ROS would trigger further activation of Ca^2+^ channel(s), leading to the full activation of Ca^2+^ signaling and Ca^2+^-based regulation of RBOHD. Furthermore, CRK2 binds the C-terminal domain of RBOHD and phosphorylates it on several specific sites (4). The right figure shows the PB1CP-mediated RBOHD down-regulation model during PTI. Upon PAMP perception, PB1CP competes with BIK1 for the binding with the N-terminal domain of RBOHD (1). PB1CP also induces relocalization of RBOHD to the endomembrane compartments (2).

